# Club cell TRPV4 as a damage sensor driving lung allergic inflammation

**DOI:** 10.1101/773382

**Authors:** Darin L. Wiesner, Richard M. Merkhofer, Carole Ober, Greg C. Kujoth, James E. Gern, Rebeca Brockman Schneider, Michael D. Evans, Daniel J. Jackson, Thomas Warner, Nizar N. Jarjour, Stephane J. Esnault, Michael B. Feldman, Matthew Freeman, Hongmei Mou, Jatin M. Vyas, Bruce S. Klein

## Abstract

Airway epithelium is the first body surface to contact inhaled irritants and report danger. We studied how epithelial cells recognize and respond to protease, which is a critical component of many allergens that provoke asthma. In a murine model, the aeroallergen alkaline protease 1 (Alp1) of *Aspergillus sp.* elicited helper T (Th) cell-dependent lung eosinophilia. Bronchiolar club cells responded rapidly to Alp1 by coordinating the accumulation of allergic immune cells in the lung. Alp1 degraded bronchiolar cell junctions, and club cells within the bronchioles propagated this signal via calcium and calcineurin to incite inflammation. In two human cohorts, we linked fungal sensitization and asthma with SNP/protein expression of the mechanosensitive calcium channel, TRPV4. TRPV4 was also necessary and sufficient for club cells to sensitize mice to Alp1. Thus, club cells detect junction damage as mechanical stress, which signals danger via TRPV4, calcium and calcineurin to initiate Th cell sensitization.

**Graphical Abstract:** 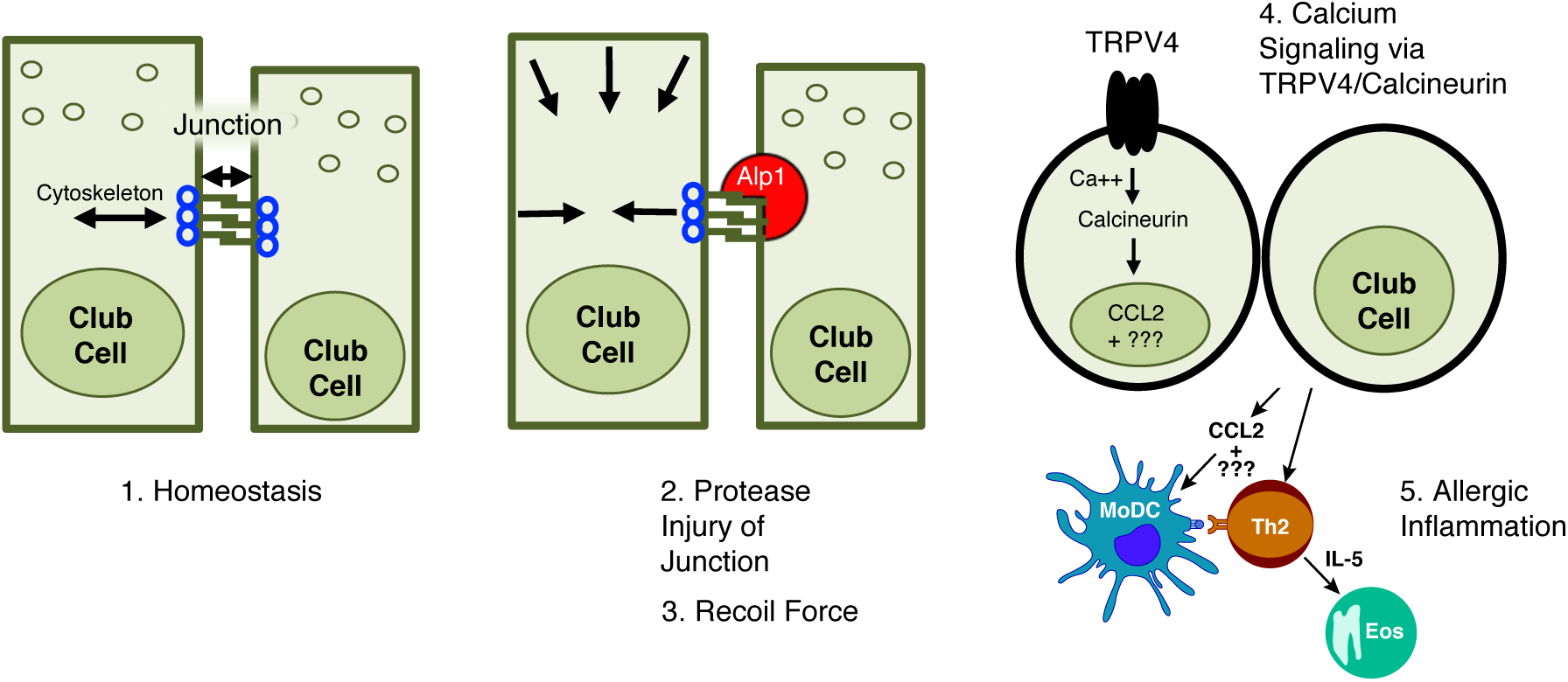

## INTRODUCTION

Asthma is often triggered by inhalation of environmental allergens, many produced by household molds (Denning et al., 2006; Denning et al., 2013; Eggleston, 2009; Hogan and Denning, 2011; Knutsen et al., 2012). *Aspergillus* is a major source of allergens *(Simon-Nobbe et al., 2008),* and alkaline protease 1 (Alp1) is the most abundant secreted protein by this mold (Sriranganadane et al., 2010; Wartenberg et al., 2011). Alp1 is a clinically important human allergen (*Asp f* 13), and the presence of Alp1 in the lungs is associated with severe asthma (Basu et al., 2018). Alp1 reportedly interrupts the interactions between smooth muscle cells and matrix components in the lung. While these events impact airway hyperreactivity, the effect of Alp1 inhalation on allergic sensitization at the lung mucosa is poorly understood.

The immune consequences of allergen exposure are well established. Briefly, type-2 helper T (Th2) cells drive IgE antibody class-switching by B cells (Lambrecht and Hammad, 2015). Th2 cells, in collaboration with innate lymphocytes (ILC), also produce cytokines that propel granulocyte recruitment, mucous production, and bronchiolar constriction (McKenzie, 2014). In contrast, the earliest events that prime this allergic cascade are just beginning to be appreciated (von Moltke and Pepper, 2018). The lung epithelium interfaces with the host and allergen and functions as both a mechanical barrier and dynamic responder (Wiesner and Klein, 2017). Upon allergen exposure, lung epithelial cells rapidly release signals that lead to type-2 leukocyte accumulation in the lungs (Roy et al., 2012; Van Dyken et al., 2014). However, the lung epithelium is not a uniform tissue, and a lack of appreciation for the heterogeneity in the epithelium has impeded our understanding of how epithelial cells recognize and respond to allergens (Wiesner and Klein, 2017).

Type-2 immune responses, besides promoting allergies, have a beneficial role in wound repair (Gause et al., 2013). Many allergens are proteases, which suggests that allergic diseases may arise when proteolytic damage to the airway is followed by dysregulated wound healing (Holgate, 2007). In fact, airway injury and loss of barrier function is a correlate of allergic disease in humans (Bousquet et al., 2000). However, the mechanisms by which epithelial cell barrier damage leads to Th cell sensitization represents a major gap in our knowledge. Airway integrity is maintained by junction proteins that mechanically link adjoining epithelial cells, and intercellular tension is balanced by intracellular forces exerted through the cytoskeleton (Ng et al., 2014). These forces are tightly regulated, and mechanosensing at the junction governs epithelial morphogenesis and cytokinesis (Pinheiro and Bellaiotache, 2018). We explored the possibility that protease damage to the junction causes the epithelium to experience a mechanical recoil force that initiates proinflammatory signaling.

Transient receptor potential (TRP) channels are a family of proteins that sense an array of stimuli, including chemicals, cold, pain, light, and pressure (Venkatachalam and Montell, 2007). To understand how the epithelium may sense mechanical strain, we investigated a particular TRP channel (i.e. TRPV4) that has osmosensory (Liedtke et al., 2000; Strotmann et al., 2000) and mechanosensory functions in various tissues (Lyons et al., 2017; Mendoza et al., 2010; O’Conor et al., 2014; Suzuki et al., 2003; Yin and Kuebler, 2010). TRPV4 is a gated calcium channel, and calcium ion currents are rapid signaling events that can occur when a cell is perturbed (Shannon et al., 2017; Zhao et al., 2006). This signaling pathway is best studied in leukocytes where calcineurin is a calcium-responsive phosphatase that regulates the activity of nuclear factor of activated T cells (NFAT) (Crabtree and Olson, 2002), which in turn transcribes a variety of proinflammatory genes.

Herein, we exploited Alp1 to test the hypothesis that allergen protease is initially sensed as damage, and this signal is translated as mechanical recoil detected via TRPV4 in epithelial cells. We identify an epithelial cell subset in the bronchioles - club cells - that responds to junction damage and elicits calcineurin-dependent inflammation. We further show that *TRPV4* is linked to fungal sensitization and asthma in humans, and is highly expressed in the bronchioles of human asthmatic patients. Finally, we establish a causal relationship between club cell TRPV4 and allergic inflammation in our fungal asthma murine model. Collectively, our data indicate that mechanical stress due to protease erosion of the epithelial cell junction creates a danger signal sensed by TRPV4 on bronchiolar club cells, which initiates a rapid calcium flux that evokes calcineurin-driven allergic sensitization.

## RESULTS

### Inhalation of *Aspergillus* alkaline protease 1 induces helper T cell-dependent eosinophilia

*Aspergillus fumigatus* is a ubiquitous household mold that abundantly secretes a human aeroallergen, alkaline protease 1 (Alp1, *Asp f*13) (Namvar et al., 2015). We used a *Pichia pastoris* expression system to produce recombinant Alp1. A ∼32 kDa protein corresponding to the size of mature Alp1 was present in the culture supernate of the Alp1-expressing strain of *Pichia* and not in the untransformed control **(Sup. Fig. 1A)**. Likewise, strong proteolytic activity was detected in supernate collected from the recombinant strain and absent when the sample was heat-inactivated or in the supernate of the untransformed control **(Sup. Fig. 1B)**.

We established a murine model to investigate the immunologic events that follow inhalation of Alp1 (**Fig. 1A**). In contrast to heat-inactive Alp1, repeated instillation of enzymatically active protease caused allergic pathology in the bronchiolar region of the lung by day 15, including leukocyte infiltration (H&E) and goblet cell metaplasia and hyperplasia (PAS) (**Fig. 1B**). We quantified the lung cellular immune response to protease by flow cytometry, excluding cells in the blood vasculature from analysis (Anderson et al., 2014). Cell suspensions from lung digests were stained with myeloid and lymphoid antibody panels with overlapping fluorophores, which allowed simultaneous quantification of 15 leukocyte- and functional-subsets that accounted for >90% of the total CD45+ hematopoietic cells in the lungs **(Sup. Fig 2A-C)**. We observed that eosinophils increased ∼500-fold (relative to heat-inactivated protease) and represented the dominant leukocyte subset by 15 days post-inoculation (dpi), outnumbering all other immune cells combined in the lungs **(Fig. 1C, Sup. Fig 1C)**.

**Figure 1.**
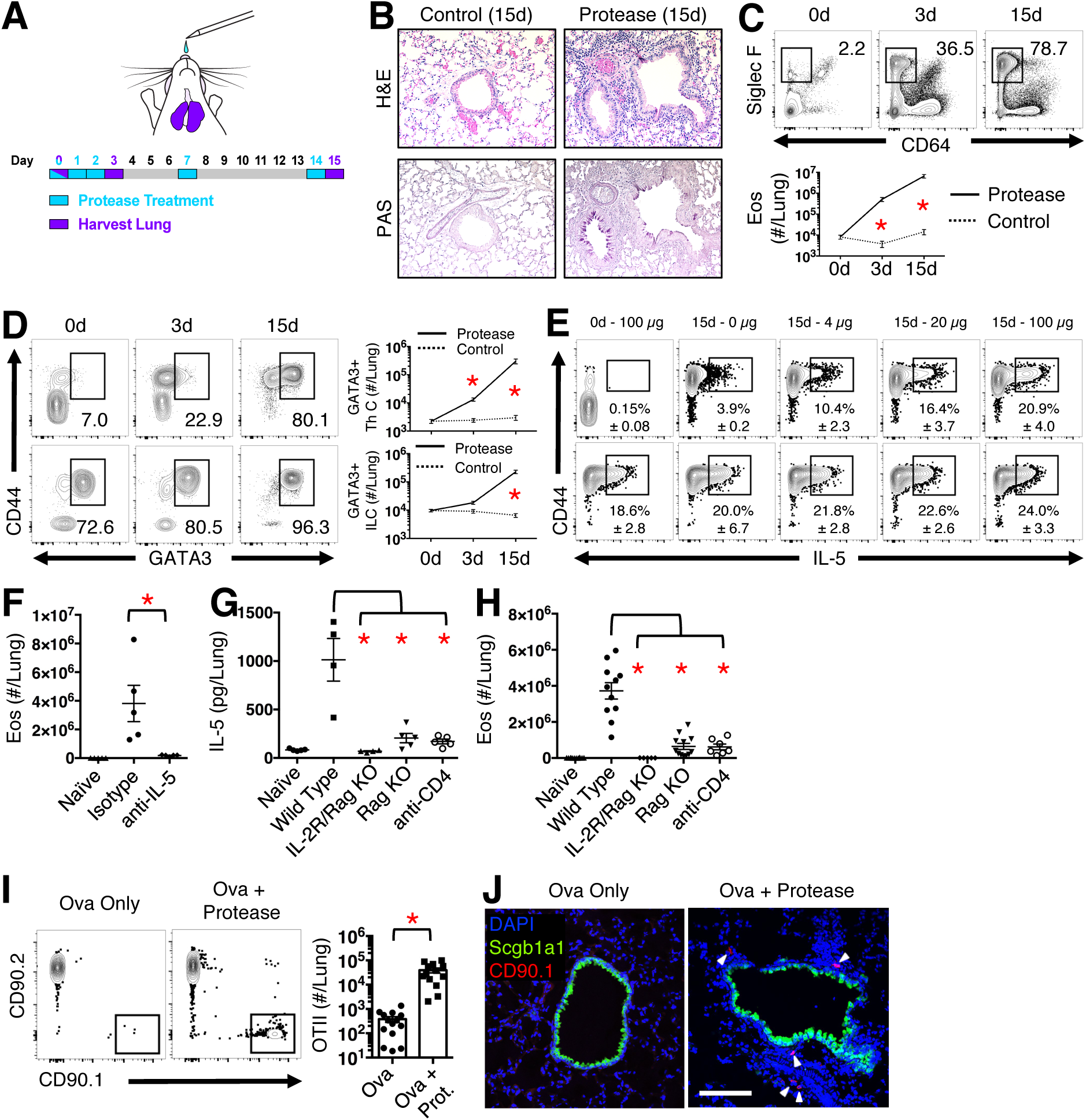
Inhalation of *Aspergillus* alkaline protease 1 induces helper T cell-dependent eosinophilia. **A)** Protease treatment schedule. **B)** Serial lung sections indicating leukocyte infiltration (H&E) and mucous production (PAS) of conducting airways of mice 15 dpi with protease or control (heat-inactive protease). **C)** Siglec F+ CD64-eosinophil accumulation in the lungs of protease treated mice. **D)** ThC (top) and ILC (bottom) expression of GATA3 in response to protease. **E)** IL-5 production by ThC (top) or ILC (bottom) upon *ex vivo* restimulation with indicated quantities of heat-inactivated protease. **F)** Eosinophils in the lungs of mice 15 dpi (or naïve) after treatment with anti-IL-5 antibody (or isotype control) **G)** IL-5 or **H)** eosinophil quantification from lungs of protease-treated (or heat inactive control), wild type and knockout animals at 15 dpi. Flow plots **I)** or immunofluorescence - 20X objective **J)** of CD90.1 OTII T cells (arrow heads) in lungs of mice 15 dpi. **Statistics**: unpaired, Mann-Whitney U test with Bonferroni adjustment for multiple comparisons; * *P* < 0.05. **Abbreviations**: dpi = days post-inoculation, Eos = Eosinophils, H&E = Hematoxylin/Eosin, IL = Interleukin, ILC = Innate Lymphoid Cell, PAS = Periodic acid-Schiff, Rag = Recombination activating gene, ThC = T helper cell. Panels B and J are representative images of at least two independent experiments. Other panels provide compiled data of two or more independent experiments.

Eosinophil differentiation is regulated by Interleukin (IL)-5, which is produced mainly by lymphocytes in non-lymphoid tissues and transcriptionally regulated by GATA3 (Lambrecht and Hammad, 2015; Van Dyken et al., 2016). We analyzed the kinetics of GATA3 and IL-5 expression in CD4+ helper T (Th) cells and innate lymphoid cells (ILC) in the lungs of protease-challenged mice to determine which of these lymphocytes is involved in the eosinophilia. Alp1 exposure resulted in expansion of GATA3^+^ Th cells and ILC in the lungs (**Fig. 1D**). Pulmonary leukocytes from unchallenged and challenged mice were stimulated *ex vivo* with titrations of heat-inactive protease to assess the capacity of Th cells and ILC to produce IL-5. Th cells from challenged mice produced IL-5 in an antigen dose-dependent manner, whereas ILC were less responsive to Alp1 (**Fig. 1E**). IL-5 neutralization abrogated the eosinophilic response to protease, indicating IL-5 is an absolute requirement for eosinophilia (**Fig. 1F**). Complete lymphocyte deficiency in IL-2Rg/Rag1^-/-^ mice resulted in a failure to produce IL-5 or recruit eosinophils (**Fig. 1G&H**). Elimination of only Th cells (with CD4 antibody or Rag deficiency) reduced both IL-5 and eosinophil accumulation to naïve levels (**Fig. 1G&H**).

We also adoptively transferred CD90.1+ ovalbumin-specific T (OTII) cells into mice to track the anatomical location of antigen-specific cells in the lungs. OTII cells accumulated in the bronchiolar region of the lung when both ovalbumin and protease were present in the inoculum (**Fig. 1I&J**). Thus, in this model, allergen-specific, GATA3+ Th cells that accumulate around the bronchioles produce IL-5 to drive a hallmark feature of allergic asthma, eosinophilia, and this allergic response requires enzymatically active Alp1.

### Bronchiolar club cells recruit monocyte-derived dendritic cells to the lungs and sensitize allergic response to Alp1

We sought to identify an antigen presenting cell population in the lungs that directs Th cell responses to fungal protease. MHCII+ CD11c+ CD11b+ CD64+ monocyte-derived dendritic cells (Mo-DC) rapidly accumulated in the lungs within 3 dpi (**Fig. 2A**). Upon pulmonary instillation of an epitope-tagged fluorescent protein (EαRFP) at 3 dpi, more than a third of the Mo-DC captured fluorescent antigen, and presented Eα peptide on MHCII (**Fig. 2B**). C-C chemokine receptor 2 (CCR2) is critical for monocyte recruitment from the bone marrow and Mo-DC formation in the lung (Leon et al., 2005). To test the requirement of Mo-DC for Th cell responses to Alp1, we transferred OTII cells into wild type and CCR2^-/-^ mice and treated the mice with ovalbumin alone or together with protease. ∼50-fold more OTII cells accumulated in the lungs of wild type mice inoculated with protease and ovalbumin than wild type mice treated with ovalbumin alone or CCR2^-/-^ mice treated with protease and ovalbumin (**Fig. 2C**). Thus, Mo-DC arrive in the lung early in the response to protease, capture and present antigen, and promote pulmonary accumulation of Th cells.

**Figure 2.**
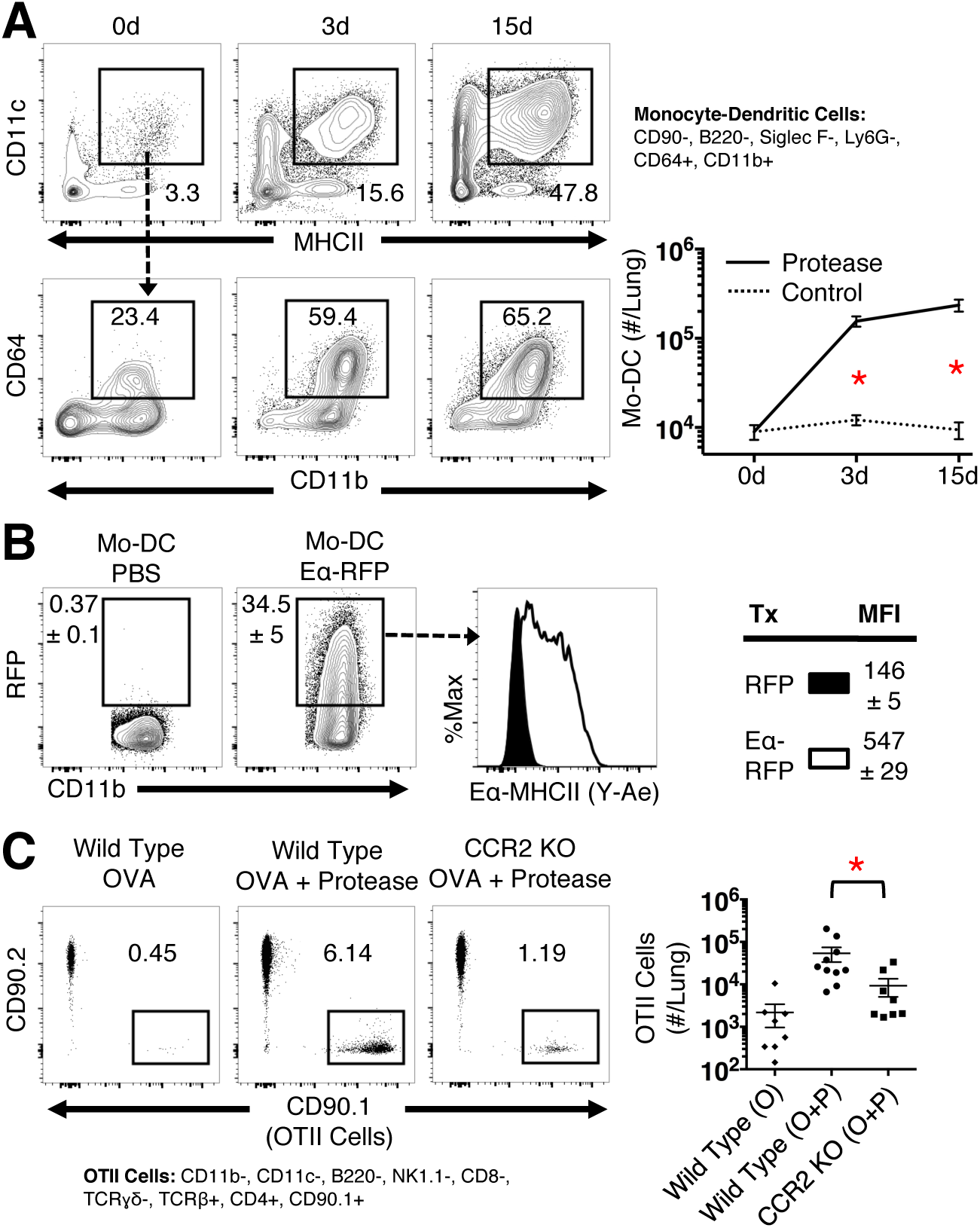
Monocyte-derived dendritic cells promote pulmonary helper T cell accumulation in response to fungal protease. **A)** Kinetic response of CD11c+ MHCII+ CD64+ CD11b+ Mo-DC after protease challenge. **B)** Mo-DC antigen uptake (Top) and presentation on MHCII (bottom) 6 hours post-inhalation of PBS, RFP, or Eα-RFP. **C)** CD90.2 endogenous and CD90.1 OTII helper T cells from lungs of wild type and CCR2^-/-^ mice 15d post-treatment with indicated antigens. **Statistics**: unpaired, Mann-Whitney U test with Bonferroni adjustment for multiple comparisons; * *P* < 0.05. **Abbreviations**: CCR = C-C Chemokine Receptor, dpi = days post-inoculation, Eos = Eosinophils, IL = Interleukin, MHC = Major Histocompatibility Class, MFI = Mean Fluorescence Intensity, Mo-DC = Monocyte-derived Dendritic Cells, Ova = Ovalbumin, RFP = Red Fluorescent Protein. Panels B and J are representative images of at least two independent experiments. All panels provide compiled data of two or more independent experiments.

C-C chemokine ligand (CCL) 2 is one ligand for CCR2 (Deshmane et al., 2009). CCL2-deficiency had a more modest effect on Mo-DC recruitment, Th2 cell accumulation, and eosinophilia after Alp1 exposure than did CCR2-deficiency **(Sup Fig. 3A&B)**, but these responses were still significantly reduced in CCL2^-/-^ mice vs. wild type controls **(Sup Fig. 3B)**. Since CCL2 significantly affects Mo-DC accumulation in the lungs, we investigated the initial source of CCL2 in the lungs of Alp1-inoculated mice. Three types of cells could produce CCL2 in response to the protease: resident phagocytes, resident lymphocytes, or stromal tissues. To determine which population is required, we ablated phagocytes with clodrosomes **(Sup Fig. 3C)**, genetically eliminated lymphocytes (IL-2Rg/Rag^-/-^), or did both. Within 8 hours of Alp1 inhalation, CCL2 levels were several fold higher in wild type mice than naive controls (**Fig. 3A**). The CCL2 response to protease remained at or above wild type mouse levels even in the absence of phagocytes and/or lymphocytes, indicating that stromal tissues are an important early source of CCL2 in this model (**Fig. 3A**).

**Figure 3.**
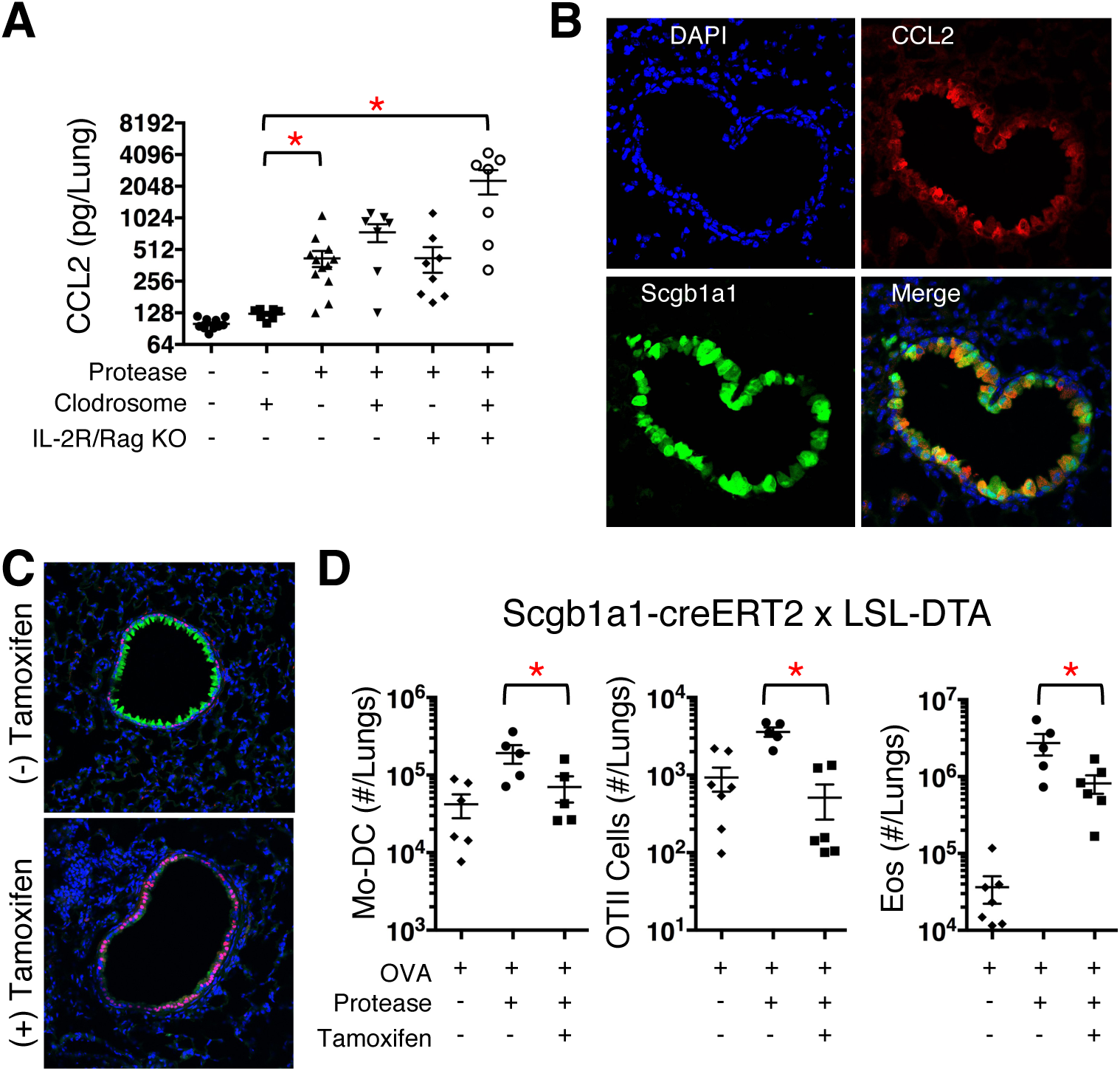
Bronchiolar club cells recruit monocyte-derived dendritic cells to the lungs and sensitize allergic response to fungal protease. **A)** CCL2 measured in lung homogenates of protease- and control-treated wild type or lymphocyte-deficienct mice. A subgroup of these mice received clodrosomes to deplete phagocytes in the lungs. **B)** Immunofluorescence images of club cells (Scgb1a1+) and CCL2 from mice treated with protease 6 hours prior to harvest. **C)** Scgb1a1-creERT2 x LSL *Diptheria* Toxin mice +/- tamoxifen every day for 4 days. Green = Scgb1a1, Red = Foxj1, Blue = DAPI. 200X magnification. **D)** Enumeration of Mo-DC, OTII T cells, and eosinophils from lungs of club-cell depleted mice and phenotypically wild mice. **Statistics**: unpaired, Mann-Whitney U test with Bonferroni adjustment for multiple comparisons; * *P* < 0.05. **Abbreviations**: CCL = Chemokine Ligand, CCR = Chemokine Receptor, dpi = days post-inoculation, Eos = Eosinophils, IL = Interleukin, Mo-DC = Monocyte-derived Dendritic Cells, Ova = Ovalbumin, Scgb1a1 = Secretoglobin. Panels B and C are representative images of at least two independent experiments. Other panels provide compiled data of two or more independent experiments.

Epithelial cells release CCL2 in response to sterile lung injury and inhaled chitin (Mercer et al., 2009; Roy et al., 2012). However, the lung epithelium is not a homogenous tissue and is comprised of at least seven distinct subsets with varying functions (Wiesner and Klein, 2017). We use confocal microscopy to identify if/which epithelial cell subset produces CCL2. Within hours of Alp1 exposure, CCL2 expression was confined to secretoglobin (Scgb1a1)+ club cells in the bronchiolar regions of the lung **(Fig. 3B, Sup Fig. 3D)**. In mice with Scgb1a1-estrogen receptor cre and cre-inducible *Diptheria* toxin, tamoxifen administration eliminated club cells, which were replaced by Forkhead box J1 (Foxj1)+ ciliated epithelial cells (**Fig. 3C**). This system allowed us to test the requirement of club cells in response to protease. In Club cell-depleted mice the numbers of Mo-DC and OTII cells in the lung returned toward baseline, and eosinophil recruitment fell sharply by 15 dpi (**Fig. 3D**). Thus, an individual epithelial cell subset (i.e. club cells) is a driver of allergic inflammation. Interestingly, club cells are a major constituent of the bronchiolar epithelium, which coincides with the focal areas of inflammation in **Figure 1B&J**.

### Proposed models that fail to explain Alp1 instigated allergic inflammation

To understand how protease is recognized by the epithelium, we tested several models. Investigators have proposed that fungal proteases cleave host proteins, coopting normally physiologic pathways to cause atopic disease (Cayrol et al., 2018; Drouin et al., 2002; Kauffman et al., 2000; Millien et al., 2013). We tested the roles of (i) protease activated receptors (PAR) 1 or 2, (ii) complement C3, (iii) fibrinogen, and (iv) IL-33, as sensors of Alp1. First, Alp1 failed to cleave at the precise site of PAR1 or PAR2 required for tethered-ligand signaling **(Sup. Fig. 4A&B)**. Likewise, the allergic response to Alp1 in whole genome PAR1- or PAR2-knockout mice was not different from wild type animals **(Sup. Fig. 4C&D)**. Second, we asked if Alp1 cleaves complement C3 to generate C3a anaphylatoxin, which could bind C3a receptors on club cells (Drouin et al., 2002). We found that club cells display C3a receptor, and that Alp1 excises a fragment from C3 similar in size to C3a **(Sup. Fig. 5A&B)**. However, MALDI-TOF analysis revealed that the resultant fragment was not identical in sequence to C3a, and the allergic response to Alp1 was also unaffected in C3-deficient mice **(Sup. Fig. 5C&D)**.

We next tested whether Alp1 releases fibrinopeptides from fibrinogen to signal inflammation. We found that the largest amount of inhaled thrombin - the natural enzyme for fibrinogen - that mice could survive (2000 units), had only a subtle effect on inflammation and was not additive or synergistic with Alp1 **(Sup. Fig. 6A)**. Likewise, hirudin, a thrombin inhibitor, failed to suppress inflammation 3dpi when administered together with Alp1 **(Sup. Fig. 6B)**. Alp1 neither induced clot formation when incubated with human fibrinogen, nor released fibrinopeptides A or B into the reaction mixture **(Sup. Fig. 6C&D).** Furthermore, the reaction mixture was only able to induce inflammation if the sample was not heat-inactivated to remove protease activity **(Sup. Fig. 6E).** Lastly, we found that the presumptive receptor for fibrinogen cleavage products, toll-like receptors (Millien et al., 2013), were dispensable for the allergic response to Alp1 **(Sup. Fig. 6F)**. MyD88, the downstream signaling intermediate for both toll-like receptors and IL-33 receptor, also was not required in this model of allergic inflammation **(Sup. Fig. 6F).** Of note, ILC2 numbers were reduced to naive levels in protease-treated MyD88^-/-^ mice, suggesting that the inflammatory effects of IL-33 involve ILC2 cells more than Th2 cells **(Sup. Fig. 6F)**.

### Protease damage to epithelial junctions in the bronchioles instigates inflammation

Alp1 lacks an exosite that would render it specific for other mammalian proteins beyond those mentioned in the preceding paragraph. Therefore, we modified our hypothesis and tested if fungal protease broadly disrupts the epithelial barrier. Inhalation of biotin in the presence or absence of Alp1, followed 1 hour later by lung processing for microscopy, revealed that the protease caused biotin to leak from the bronchiolar regions of the lung, indicating a loss of airway integrity (**Fig. 4A**).

**Figure 4.**
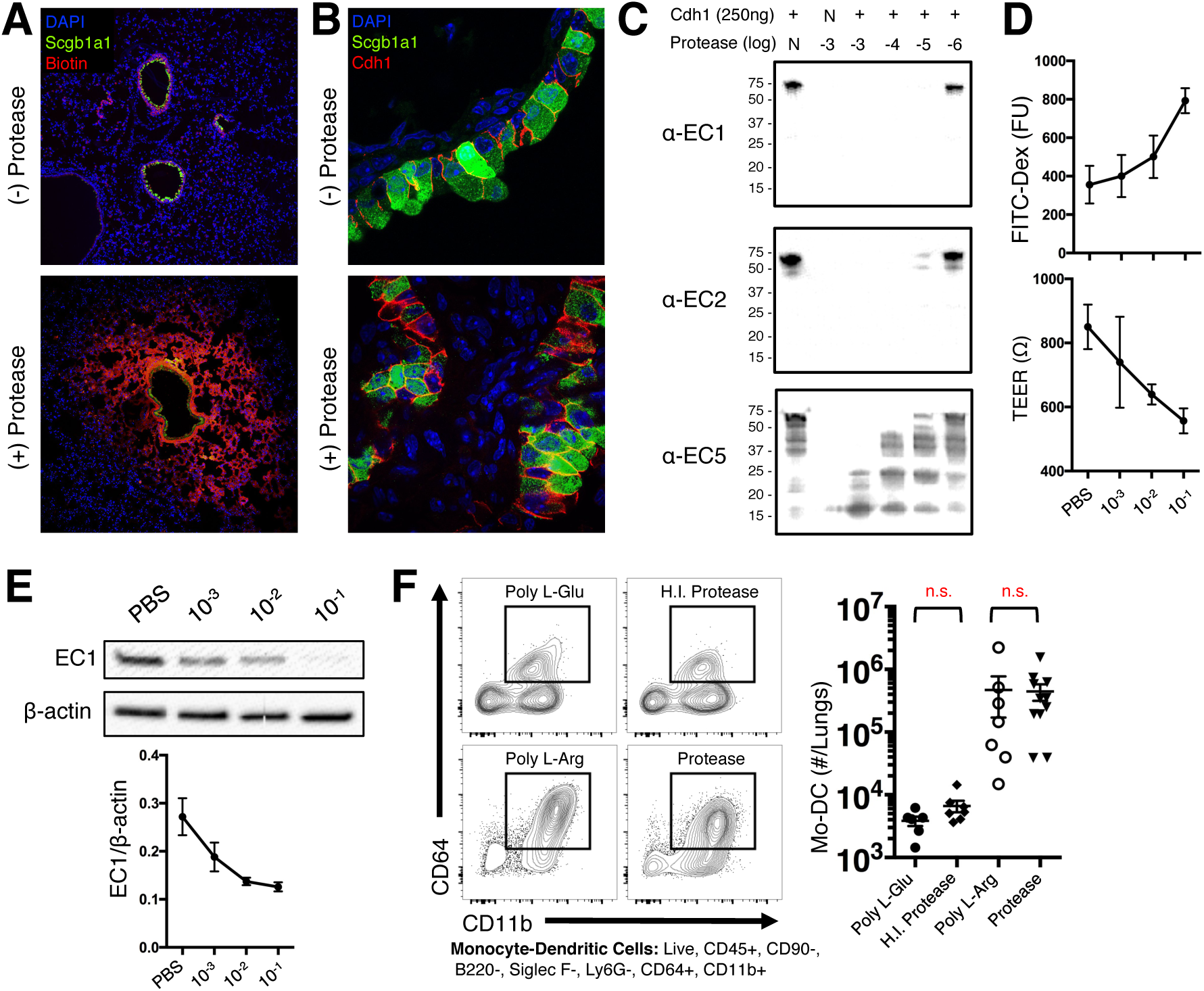
Protease damage to epithelial junctions in the bronchioles instigates inflammation. **A)** Biotin leakage from the airways with (bottom) or without (Top) fungal protease 1H post-treatment. 20X objective. **B)** Immunofluorescence of lungs 3 dpi. 100X objective. **C)** Purified Cdh1 incubated with log-fold dilutions of protease and blots probed with monoclonal antibodies targeting Cdh1 extracellular domains. **D)** Primary human lung epithelial cells grown in an air-liquid interface incubated with log dilution of protease. Barrier integrity was quantified by FITC-dextran leakage or TEER. **E)** Lysates from panel D were blotted for EC1. **F)** Mice received 25μg of either Poly-L-Glu, Heat-inactive Protease, Poly-L-Arg, or Protease for 3 consecutive days. Lungs were processed for flow cytometry, and the Mo-DC response was quantified. **Statistics**: unpaired, Mann-Whitney U test with Bonferroni adjustment for multiple comparisons; * *P* < 0.05. **Abbreviations**: Arg = Arginine, CCL = Chemokine Ligand, CCR = Chemokine Receptor, Cdh1 = E-cadherin, Dex = Dextran, dpi = Days Post-inoculation, EC = Extracellular, Eos = Eosinophils, Glu = Glutamate, IL = Interleukin, Mo-DC = Monocyte-derived Dendritic Cells, Ova = Ovalbumin, Scgb1a1 = Secretoglobin, TEER = Transepithelial Electrical Resistance. Panels A, B and C are representative images of at least two independent experiments. Other panels provide compiled data of two or more independent experiments.

The mechanical barrier of the airway is maintained by solute impermeable tight junctions (TJ) at the apical reach of the epithelia, while adherens junctions (AJ) are situated just beneath the TJ. Whereas TJ proteins are heterogeneous in their identity and expression pattern, the AJ is formed through highly conserved, homotypic interactions of E-cadherin between adjacent epithelial cells (Hartsock and Nelson, 2008; Perez-Moreno and Fuchs, 2006). We posited that measuring the effect of Alp1 on E-cadherin is a direct way to test whether protease significantly disrupts the epithelial junction. Commercial antibody to mouse E-cadherin (clone DECMA-1) recognizes the cytosolic portion of E-cadherin, which appeared unaffected by Alp1 in the bronchioles of mice despite the hyperplastic appearance of the airway cells (**Fig. 4B**).

Some monoclonal antibodies recognize individual extracellular (EC) domains of E-cadherin (Shiraishi et al., 2005). To test if Alp1 degrades E-cadherin in the absence of confounding host enzymes (Zemans et al., 2011), we incubated recombinant E-cadherin with titrations of Alp1 for 1 hour. Western blots showed that even low concentrations of Alp1 destroyed the epitopes of E-cadherin recognized by antibodies against EC1 and EC2 (**Fig. 4C**). Next, we grew human bronchiolar epithelial cells at an air-liquid interface and tested whether Alp1 penetrates the TJ and damages E-cadherin *in situ*. After 1 hour, Alp1 compromised barrier integrity (**Fig. 4D**), as measured by FITC leakage and reduced transepithelial electrical resistance (TEER), and also degraded the EC1 domain of E-cadherin (**Fig. 4E**).

Cationic polypeptides such as poly-L arginine interrupt epithelial junctions (Shahana et al., 2002). When this compound was administered into the lungs of mice, direct junction damage (without the intrinsic ability to cleave host proteins) was sufficient to drive Mo-DC accumulation 3 dpi (**Fig 4F**). Collectively, these data establish that Alp1 disrupts epithelial junctions, and that damage to junctions between conducting airway cells signals inflammation.

### Allergic response to fungal protease requires calcineurin signaling in club cells

We investigated the signaling events in club cells that initiate inflammation upon protease injury to the epithelial junction. The AJ is at the center of a critical signaling pathway involved in spatial patterning during lung development, cancer metastasis, and epithelial wound healing (Perez-Moreno and Fuchs, 2006). Many of these processes feed into the Wnt pathway, where β-catenin dissociates from E-cadherin at the AJ and migrates to the nucleus (Komiya and Habas, 2008). Although active β-catenin was expressed at the lateral membrane of club cells, active β-catenin did not concentrate in the nucleus of club cells in protease-damaged regions of the bronchioles (**Fig. 5A**). Nuclear Factor κ Light Chain of B cells (NFκB) is another major signaling pathway implicated in epithelial cell responses to allergen (Lambrecht and Hammad, 2014; Whitsett and Alenghat, 2015). However, protease-challenged mice with pan-lung epithelial cell deficiency in the positive regulator of NF-κB, inhibitor of nuclear factor kappa-B kinase subunit beta (IKK2) (Perez-Nazario et al., 2013), did not elicit GATA3+ Th cell response differently from wild type littermates (**Fig. 5B**). Thus, unexpectedly, neither Wnt nor NF-κB signaling is responsible for club cell-dependent inflammation.

**Figure 5.**
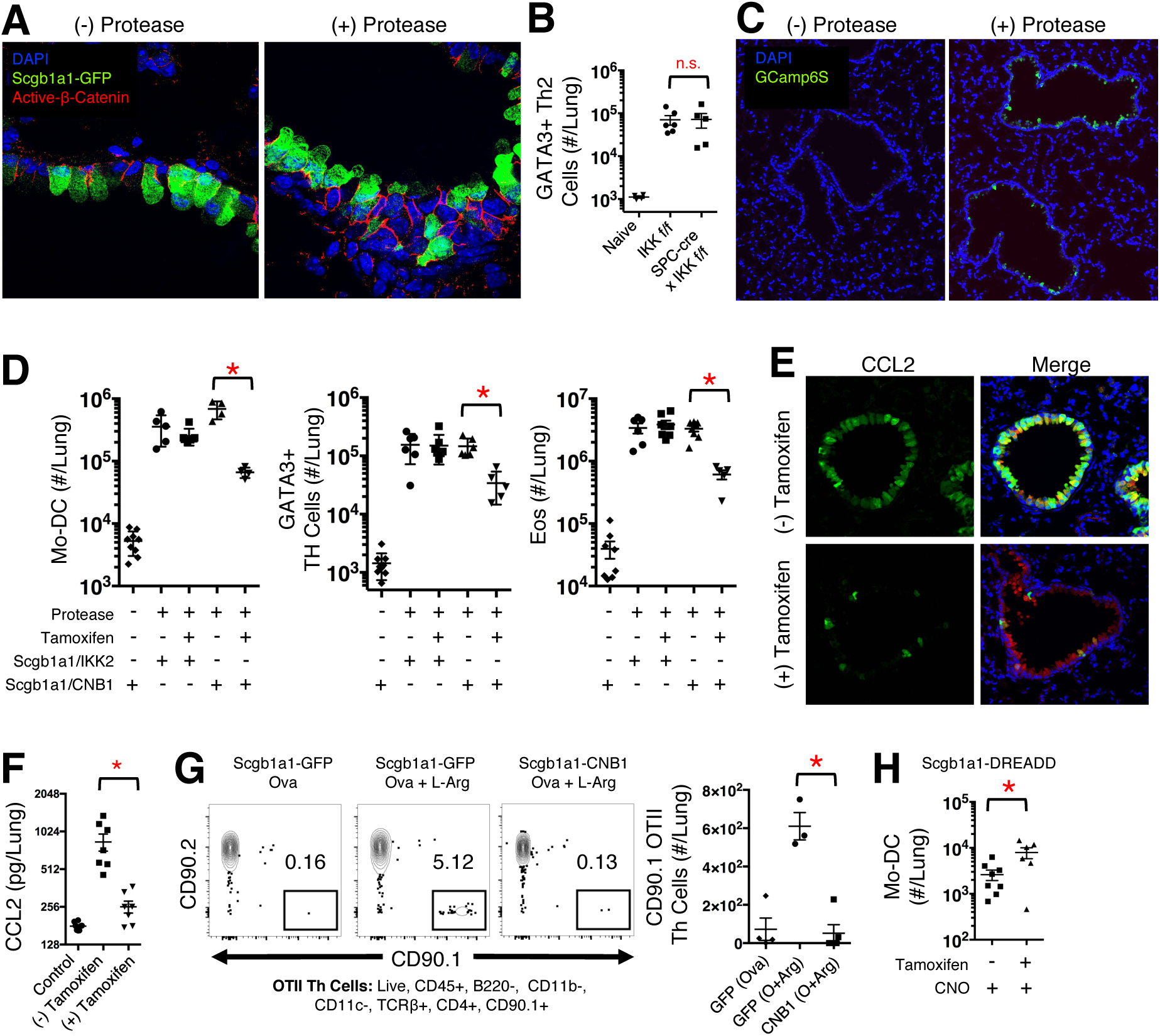
Allergic response to Alp1 requires club cell calcineurin. **A)** Immunofluorescence images of lungs 3dpi. 100X objective. **B)** GATA3+ CD4+ T cells from lungs of mice with NF-κB deficiency in entire lung epithelium 15 dpi protease exposure. **C)** Scgb1a1-creERT2 x GCamp6S (calcium indicator protein) mice treated with control (left) or protease (right). Green = Calcium flux. Blue = Nuclei. 20X Objective. **D)** Cellular response to protease 15 dpi in mice with Club Cell deficiencies in NF-κB (IKK2) or NFAT (CNB1) pathways. **E)** Immunofluorescence images collected from Scgb1a1-CreERT2 x CNB1 floxed mice treated with or without tamoxifen 3 dpi with protease. Green = CCL2, Red = Scgb1a1, Blue = DAPI. 20X objective. **F)** CCL2 quantified from lung lysates of Scgb1a1-CreERT2 x CNB1 floxed mice treated with or without tamoxifen, analyzed 3 dpi with protease. **G)** OTII response 15 dpi with Ova, with or without Poly-L-Arg in wild phenotype mice or mice with calcineurin-deficiencies in Club Cells. **H)** Mo-DC from Scgb1a1-CreERT2 x DREADD (hM3Dq) mice 3 dpi treatment with agonist (CNO). **Statistics**: unpaired, Mann-Whitney U test with Bonferroni adjustment for multiple comparisons; * *P* < 0.05. **Abbreviations**: Arg = Arginine, CNB1 - Calcineurin, CCL = Chemokine Ligand, CNO = Clozapine N-Oxide, dpi = Days Post-inoculation, Eos = Eosinophils, Mo-DC = Monocyte-derived Dendritic Cells, Ova = Ovalbumin, Scgb1a1 = Secretoglobin, Th2 = Type-2 Helper T cells. Panels A, C and E are representative images of at least two independent experiments. Other panels provide compiled data of two or more independent experiments.

The findings above suggested that an atypical pathway in club cells drives inflammation to inhaled protease. Calcium signaling is another canonical inflammatory pathway in leukocytes, and we investigated calcium signaling in club cells based on what is known in leukocytes. In transgenic mice that express a calcium indicator protein, GCamp6s, specifically in club cells, these cells fluxed calcium within 15 minutes of exposure to inhaed protease (**Fig. 5C**). To assess the downstream consequences of calcium signaling, we used transgenic mice with tamoxifen-inducible calcineurin or IKK2 deficiency in club cells. Mice with IKK2-deficient club maintained unaffected allergic immune responses to protease (**Fig. 5D**), consistent with **Figure 5B**. Conversely, mice with calcineurin-deficient club cells demonstrated sharply blunted allergic responses of Mo-DC, GATA3+ Th cells, eosinophils, and CCL2 levels in responses to protease (**Fig 5D-F**). We also investigated mice lacking calcineurin or IKK2 in other epithelial cell subsets - Sftpc+ Type-2 alveolar cells, Foxj1+ ciliated cells, or Ascl1+ pulmonary neuroendocrine cells - and found no effect of these pathways on inflammation **(Sup. Fig. 7A&B)**.

Poly L-arginine targeting of the junction caused a large expansion of OTII cells in wild type mice, and this response was reduced to control levels with an induced deletion of calcineurin in club cells (**Fig. 5G**). Thus, junction damage alone can sensitize T cells, and similar to protease, this response signals via calcineurin in club cells.

Finally, we asked if calcium signaling in club cells is sufficient to drive inflammation. Scgb1a1-creERT2 mice were crossed with hM3Dq (i.e. DREADD) mice and treated with clozapine n-oxide (CNO) to induce G-protein-mediated calcium flux in club cells. Mice receiving both CNO and tamoxifen recruited a larger population of Mo-DC to the lungs compared to mice treated only with CNO (**Fig. 5H**). In summary, protease damage to the epithelial junction elicits a calcium flux in club cells that signals through calcineurin to mediate allergic inflammation.

### Apical bronchiolar epithelium expression of TRPV4 is associated with fungal asthma

We searched for a receptor/channel that could connect protease injury at the bronchiolar junction with calcium entry into club cells. We posited that TRPV4 could exert this function because it acts as a mechanosensative gated-calcium channel. Genetic variation at the *TRPV4* locus has been associated with lung pathology in prior studies in humans. In particular, the G allele or AG+GG genotypes at rs6606743 was associated with osmotic airway hyper-responsiveness (Naumov et al., 2016) and increased risk of COPD (Zhu et al., 2009). We therefore tested for the association of genotype at rs6606743 with fungal sensitization and asthma in 211 participants in the Childhood Origins of Asthma (COAST) study. We identified significant associations with both phenotypes (**Fig. 6A-B**). This SNP resides 496 kb upstream of the transcription start site of the *TRPV4* gene, within an intron of a long non-coding RNA (lncRNA), XR_945333.1. This variant is also in linkage disequilibrium (*r*^2^ >0.8 in CEU) with three SNPs spanning about 2.5kbp, located within the lncRNA gene and overlapping an enhancer site annotated in multiple cell types, including keratinocytes (NHEK) and lung epithelial cells (A549) in the Roadmap Epigenomics project (Roadmap Epigenomics et al., 2015). To directly test whether rs6606743, or SNPs in linkage disequilibrium with it, are associated with expression of the *TRPV4* gene, we used expression quantitative locus (eQTL) data from epithelial tissues, esophagus and skin, available in the GTEx database (Consortium et al., 2017). Indeed, the G allele at rs6606743 is associated with increased expression of *TRPV4* in the epithelium of esophagus (*P* = 4.5e-5) and skin (*P* = 2.4e-7). Collectively, these data suggest that increased expression of TRVP4 is associated with a risk of sensitization to fungal allergen and asthma.

**Figure 6.**
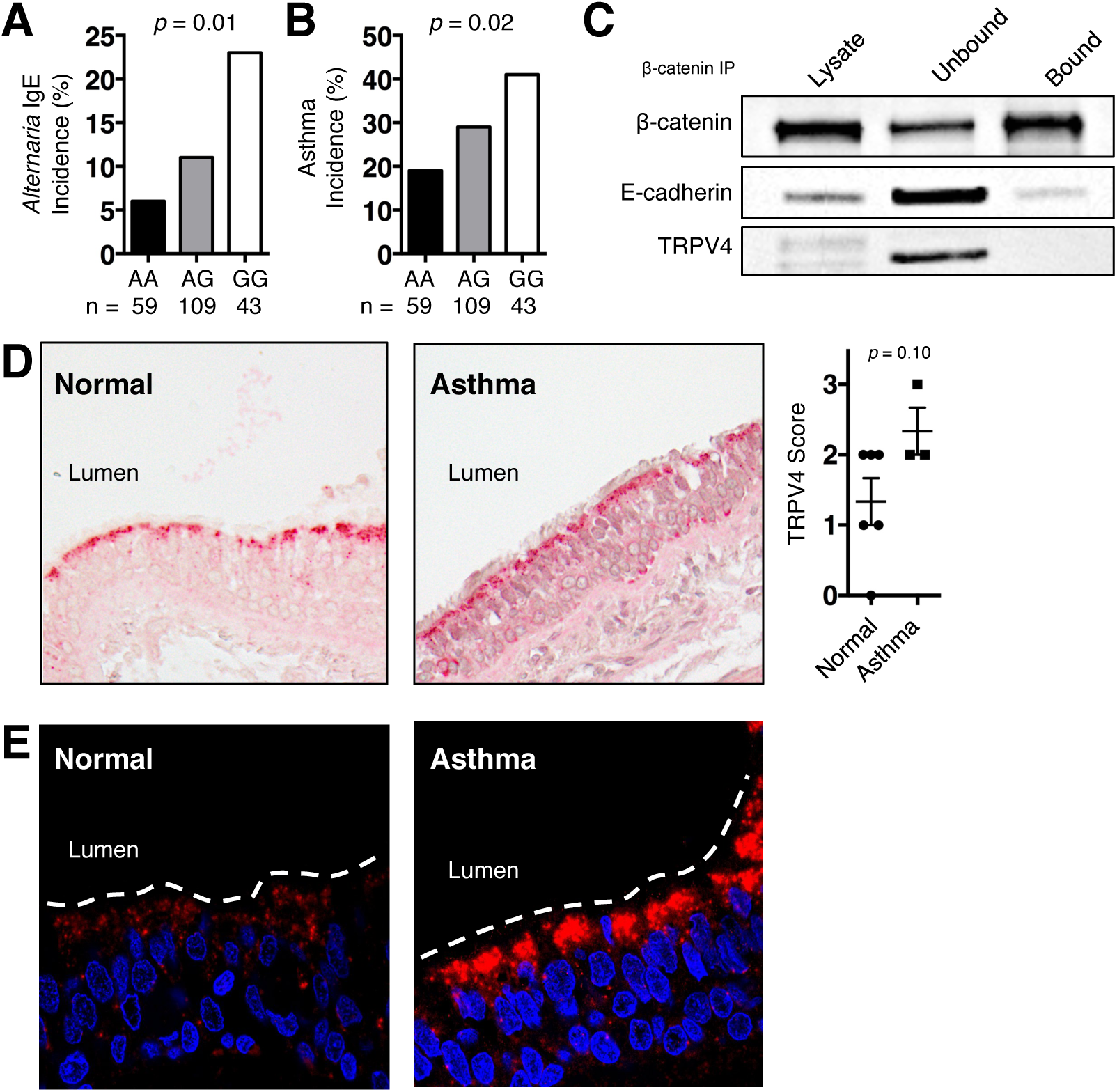
Apical membrane expression of TRPV4 is associated with fungal sensitization and asthma in humans. Chi-square test of the onset of **A)** fungal-specific IgE at 3 years and **B)** asthma 6 years associated with SNP at position rs6606743 of *TRPV4*. *N* = 211 children **C)** Western blots of lysates or membrane enriched samples (bound/unbound) subjected to immunoprecipitation with anti-β-catenin antibody from human club and goblet cells. **D)** Immmunocytochemistry (40X objective) or **E)** immunofluorescence (100X objective) of bronchial biopsies from healthy donors (*N*=5) and patients with asthma (*N*=3). TRPV4 is indicated by red chromogen and red fluorescence. Unpaired, non-parametric T test. Panels C is representative images of at least two independent experiments.

In a separate group of individuals, we analyzed the location and level of TRPV4 expression in human bronchiolar cells and biopsies. Cell lysates of club/goblet cells differentiated from primary human lung basal cells (Feldman et al., 2019) were enriched for membrane proteins (Sokabe et al., 2010). The membrane proteins were immunoprecipitated with anti-β-catenin antibody to determine if TRPV4 physically interacts with β-catenin at the AJ, as previously reported with keratinocytes (Sokabe et al., 2010). TRPV4 was more abundant in the unbound fraction compared to the lysate, indicating TRPV4 concentrates in the cell membrane (**Fig. 6C**). However, TRPV4 was absent in the bound fraction, indicating TRPV4 does not form an avid interaction with β-catenin at the AJ in human club/goblet cells (**Fig. 6C**).

We also immunologically stained bronchial biopsies for TRPV4 in 5 healthy donors and 3 patients with asthma. Intriguingly, TRPV4 was most apparent in the apical tip of the bronchiolar epithelium (**Fig. 6D&E**). TRPV4 expression was scored by a sample-blinded pathologist, and, when compared with healthy individuals, TRPV4 expression trended higher in asthmatics (**Fig. 6D**). Taken together, TRPV4 is highly expressed in the apical membrane of bronchiolar cells of asthma patients.

### Club cell TRPV4 signals allergic inflammation

To determine if a directional relationship exists between TRPV4, club cells, and Th cell sensitization, we returned to our murine model. First, we used a functional assay to assess TRPV4 expression by club cells. Cell suspensions from club cell-reporter mice were loaded with a ratiometric calcium indicator dye and analyzed by flow cytometry. Club cells rapidly shifted fluorescence in response to a TRPV4 agonist, indicating that club cells express a functional TRPV4 calcium channel (**Fig. 7A**). Second, we tested the requirement of club cell TRPV4 in protease-driven allergic inflammation by using Scgb1a1-creERT2 mice with a floxed *TRPV4* allele. Tamoxifen administration to delete *TRPV4* in club cells resulted in significantly decreased CCL2 production within hours of protease inhalation, and reduced OTII cell accumulation 15 dpi (**Fig. 7B&C**). Lastly, we modeled the human condition of enhanced TRPV4 expression by overexpressing TRPV4 *in vivo* with adeno-associated virus (AAV6). We found that AAV6 expressed (mCherry) efficiently in club cells (**Fig. 7D**). We thus infected Scgb1a1-creERT2 mice with an AAV6-doublefloxed inverse orf-TRPV4 virus that overexpresses TRPV4 in cells with a functional cre recombinase (Wheeler et al., 2016). When viral gene expression was activated by tamoxifen, Scgb1a1-creERT2 mice inoculated with protease accumulated significantly more OTII cells than wild phenotype mice not receiving tamoxifen (**Fig. 7E**). Remarkably, the enhanced OTII cell response due to TRPV4 overexpression was absent in mice with club cell calcineurin deficiency, indicating that TRPV4 and calcinueurin are in epistasis (**Fig. 7F**). In sum, these results demonstrate that club cell TRPV4 is required for CD4+ T cell sensitization to Alp1, and that recapitulating the features of human asthma by overexpressing TRPV4 enhances sensitivity to fungal protease.

**Figure 7.**
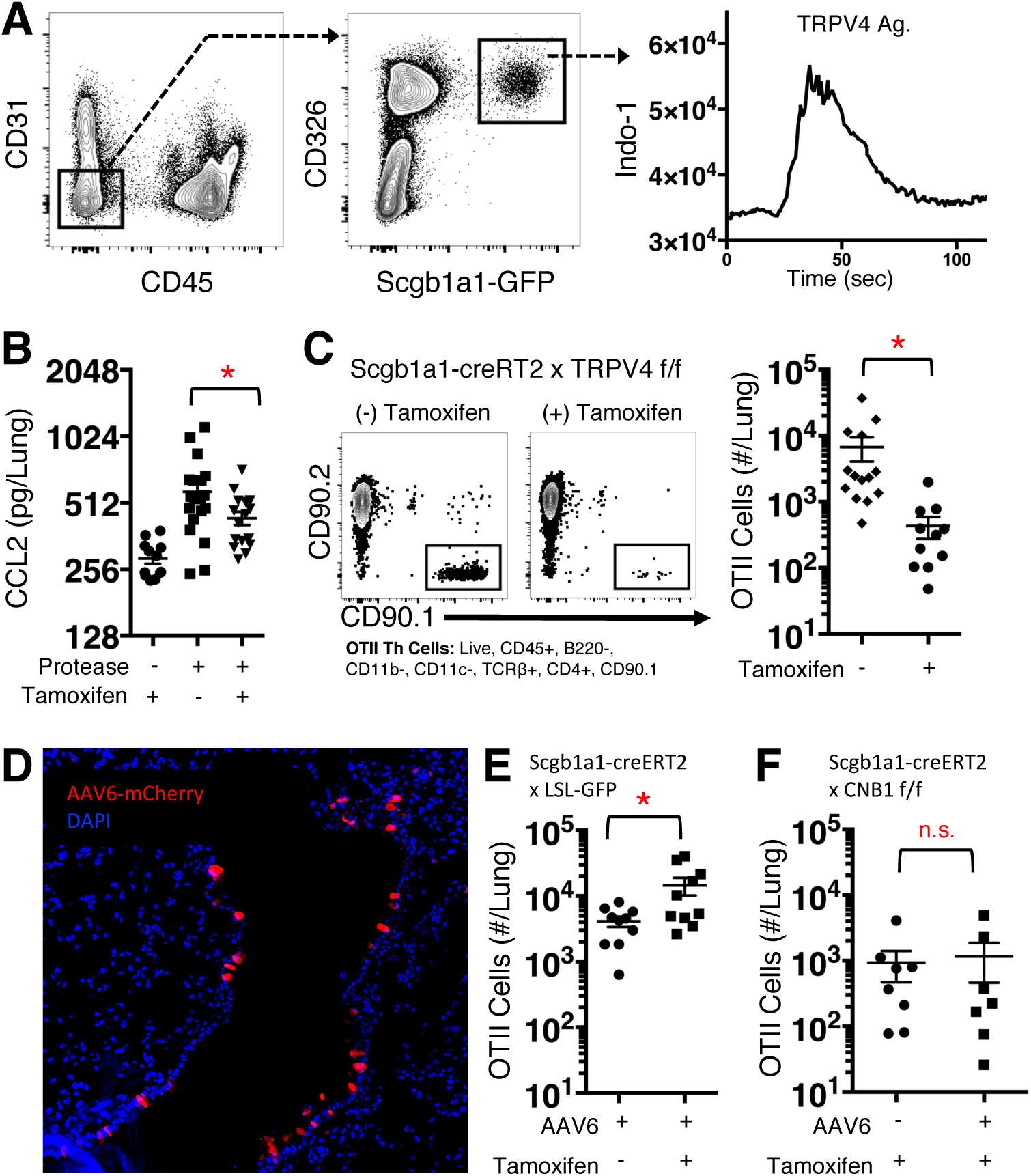
Club cell TRPV4 signals allergic inflammation. **A)** Flow plots of lung cells from Scgb1a1-GFP mice loaded with a calcium indicator dye, Indo-1, and stimulated with a TRPV4 agonist GSK1016790A. **B**) CCL2 (8hr) and **C)** CD90.1+ OTII cell (15dpi) response to protease in Scgb1a1-creERT2 x TRPV4 floxed mice. **D)** Immunofluorescent images of lung sections from mice intranasally infected with AAV6-mCherry 3 weeks prior. CD90.1 OTII T cell response to protease 15 dpi in **E)** Scgb1a1-creERT2 x LSL GFP or **F)** Scgb1a1-creERT2 x CNB1^fl/fl^ mice. **Statistics**: unpaired, Mann-Whitney U test with Bonferroni adjustment for multiple comparisons; * *P* < 0.05. **Abbreviations**: AAV = Adeno-associated virus, CCL = Chemokine Ligand,, CNB1 - Calcineurin, dpi = Days Post-inoculation, Eos = Eosinophils, LSL-GFP = Lox-Stop-Lox Green Fluorescent Protein, Mo-DC = Monocyte-derived Dendritic Cells, Ova = Ovalbumin, Scgb1a1 = Secretoglobin. Panels A and D are representative data of multiple independent experiments. Other panels provide compiled data of two or more independent experiments.

## DISCUSSION

Aeroallergens, including proteases, are irritants (Holgate, 2007), and the airway epithelium is the first tissue assaulted. We investigated how the lung epithelium alerts immune cells to protease allergen exposure, sensitizing CD4+ T cells. We report a conceptual advance in understanding the pathogenesis of asthma, revealing that the mechanical force exerted on epithelial cells that suffer junction damage is a danger signal sensed by TRPV4. We define a linear pathway connecting epithelial junction damage with calcium flux via TRPV4, calcineurin signaling, and Th cell sensitization.

Many patients with asthma are allergic to environmental molds. We chose the fungal protease Alp1 to probe the epithelium and allergic inflammation for several reasons. Alp1 is one of the most abundantly secreted *Aspergillus* proteins and a human allergen. *Aspergillus* is a saprophyte, and proteases secreted by the fungus facilitate nutrient acquisition in the environment (St Leger et al., 1997). These proteases are not virulence factors (Bergmann et al., 2009), and have not undergone host selection to evolve exosites that mammalian elastases, convertases or thrombin require for specific, avid interaction with their substrate (Overall and Blobel, 2007). Moreover, organisms spanning every reach of the biologic kingdom produce proteases that elicit allergic responses in humans (Reed and Kita, 2004). With this expansive diversity of proteases comes a broad substrate specificity that argues against a highly evolved interaction between host component(s) and protease.

We tested several possible models of protease-induced allergic inflammation. The most striking finding upon protease inhalation was damage to the bronchiolar epithelial junction. We propose a model of asthma pathogenesis in which protease injury to epithelial cell junctions elicits an unconventional “damage response” sensed as recoil by a mechanoreceptor TRPV4. This mechanism of sensing damage runs counter to classic damage receptor: ligand interactions described previously. The most widely recognized mechanism by which cells report injury is via damage-associated molecular patterns (DAMPs). In general, DAMPs are proteins (e.g. IL-33, High Mobility Group Box 1, F-actin), nucleic acids, or metabolites (e.g. ATP) that are sequestered within cells during steady state. These molecules spill into the extracellular milieu when a cell ruptures during necrosis (Kaczmarek et al., 2013). DAMPs alert a healing response in neighboring cells bearing DAMP receptors. While protease injury induced hyperplasia of the bronchiolar epithelium, we found little necrosis in the affected regions (data not shown). These observations indicate that junction disruption, independent of cell lysis/death, is a bona fide damage signal. We posit that mechanical perturbations exerted on club cells as they detach from one another are sensed by TRPV4, prompting cellular calcium flux and calcineurin-dependant inflammatory signaling. Our findings extend the known mechanosensing ability of TRPV4 (Suzuki et al., 2003), and we propose that TRPV4 moonlights as a “damage sensor” during asthma pathogenesis.

TRP proteins perform a variety of sensory functions, and several TRP channels have been implicated in asthma. TRPV4 expressed by smooth muscle cells responds to osmotic stimuli and contributes to hypotonic airway contractility (Jia et al., 2004). SNP rs6606743 of *TRPV4* has been associated with osmotic hyperresponsiveness in patients with bronchial asthma (Naumov et al., 2016). Mice with a whole genome deletion of *TRPV4* are protected from airway remodeling after exposure to house dust mite antigens (Gombedza et al., 2017). Our work extends previous studies of airway hyperreactivity and fibrosis by defining a role for epithelial cell TRPV4 in sensing damage and driving allergic inflammation. The pain sensing channel, TRPV1, and cold sensing channel, TRPM8, also expressed by lung epithelia, promote inflammatory responses when stimulated, and have been associated with asthma (Cantero-Recasens et al., 2010; Choi et al., 2018; McGarvey et al., 2014; Sabnis et al., 2008). TRPV1 and TRPM8 are calcium channels, and consequently, their involvement in epithelial responses and asthma also could be mediated by calcium signaling through calcineurin, as we found with TRPV4. Thus, calcium signaling and calcineurin might be a central pathway in the pathogenesis of allergic asthma, perhaps amenable to therapeutic targeting.

In contrast to the widely held belief that NF-κB is the master regulator of inflammatory signaling within the epithelium (Janssen-Heininger et al., 2009; Lambrecht and Hammad, 2014; Whitsett and Alenghat, 2015), we found that NF-κB had no requisite role in the epithelial response to protease. Our model is distinct in two ways that could potentially explain this discrepancy. First, we avoided LPS contamination in our allergen preparations by producing Alp1 with a yeast expression system. Many experimental allergens, like those from dust mites, are crude extracts, containing unknown quantities of LPS and other undefined contaminants (Cates et al., 2004). The presence of TLR ligands in these allergens could prompt NF-κB-dependent inflammation in epithelial cells. Second, mice only received allergen intranasally in our model, whereas other approaches, including ovalbumin/alum, require systemic sensitization followed by intranasal challenge. With intranasal sensitization, the epithelium is intimately involved in the early innate cell activation and T cell priming events, whereas challenge after systemic sensitization prompts epithelial cells to recruit activated T cells from the periphery to lung. These disparate situations may rely on a different set of signals in the epithelium.

The bronchiolar epithelium is a central feature in our model of protease driven allergic inflammation. Protease damage was concentrated in the bronchioles: (i) bronchiolar club cells had a requisite role in driving a large part of the response to protease; (ii) the inflammation was confined to bronchiolar region of the lung; and (iii) TRPV4 was highly expressed in the bronchiolar epithelium of humans. Mice and humans have similar epithelial cell subsets, and these cells retain comparable functions across species. However, the proportions of epithelial cell subsets that comprise the conducting airways (i.e. bronchioles) differ between mice and humans (Iwasaki et al., 2017). While the bronchioles are densely populated by club cells with a few intermittent ciliated cells in mice, club cells and ciliated cells are present in more balanced proportions in similar regions of human lungs. In our murine model, club cells had a potent effect, and all manipulations of the epithelium were targeted to club cells in our investigations. When we depleted club cells, Foxj1+ ciliated cells repopulated the bronchioles, and the inflammatory response to protease was reduced. Likewise, knockout of IKK or calcineurin in Foxj1+ cells did not reveal a strong phenotype in our model. Collectively, these observations suggest that ciliated cells are not sufficient or required for the response to protease. However, Foxj1+ cells expressed high levels of Ki67 after club cell depletion (data not shown), and the proliferative state of these cells could prevent them from producing proinflammatory cytokines. Also, the bronchiolar epithelium remains broadly intact when genes are disrupted in ciliated cells in mice, and club cells could perform redundant functions masking any effect that ciliated cells have in the response. Thus, the mechanisms we detailed in mouse club cells may not be restricted to only club cells in humans and could involve ciliated cells as well.

Immune responses function like electronic circuits. Cells in a network cooperate with one another to amplify a small signal into a highly ordered response. Evolution has built complexity and redundancy into the circuit, making identification of causal relationships difficult. Vaccinologists define correlates of immunity to elucidate complex biology (Plotkin, 2010). We propose that CCL2, Mo-DC, and eosinophils are correlates of immunity and not an epistatic pathway feeding into and out of Th cell sensitization. We showed that neutralizing CCL2 only modestly affects allergic inflammation, even under idealized conditions. Likewise, elimination of Mo-DC did not abrogate Th cell sensitization. Other lung resident DC populations could have redundant functions, as seen in house dust mite allergy models where CD11b+ dendritic cells overlap with Mo-DC depending on allergen dose (Plantinga et al., 2013). There are likely many signals emanating from the epithelium that instruct Mo-DC (e.g. CCL7, CCL8, CCL12, CCL13) or CD11b+ DC (e.g. Colony Stimulating Factor-1) (Moon et al., 2018). However, our goal was to understand how the epithelium recognizes and responds to protease allergen to drive Th cell sensitization. Th cell accumulation in the lungs was the primary read out in all experiments in which it was technically and biologically feasible. This criterion let us uncover a linear pathway within club cells, linking epithelial junction damage, calcium flux through TRPV4, calcineurin signaling, and Th cell sensitization.

Asthma affects nearly 10% of people in the United States and burdens the American economy in excess of $50 billion per year (Barnett and Nurmagambetov, 2011; CDC, 2013). Most cases of asthma begin in early childhood, and allergic sensitization in the first 2-3 years of life is a major risk factor for persistent childhood asthma. In our model, calcium signaling in club cells is an important mediator of Th cell priming, and elevated TRPV4 expression exacerbated the inflammatory response to protease. We also show that children with a SNP in *TRPV4* are a risk group for fungal sensitization and asthma. Genotyping at the *TRPV4* locus could identify children at increased risk for sensitization to protease-containing allergens such as fungi. These findings also suggest that targeting bronchiolar epithelial cells directly with calcineurin inhibitors could inhibit the onset of allergic sensitization, perhaps through the inhaled route to minimize systemic effects.

## ACKNOWLEDGEMENTS

We thank the following UW-Madison core facilities: University of Wisconsin Carbone Cancer Center (Grant P30 CA014520, 1S100OD018202-01), UW Biotechnology Center (NIH P50 GM64598, R33 DK070297; NSF DBI-0520825, DBI-9977525) Biochemistry Optical Core (Elle Gresvald), and Translational Research Initiatives in Pathology (Toshi Kinoshita). We Robert Gordon (Pediatrics, UW-Madison) for assistance with graphic design. Postdoctoral fellowship support was provided by the Hartwell Foundation and NHLBI T32 HL07899 (D.L.W.), and NHLBI T32 HL116275 (M.B.F.). Research grant support was provided by NIH R01 AI130411 (to B.S.K); and PO1 HL70381 and UL1 TR000427 (D.J.J., J.G., R.B.S., and C.O. in support of COAST [Childhood origins of Asthma]); R01AI136529 (J.M.V.); and Cystic Fibrosis Foundation Research Grant MOU19G0 and Charles H. Hood Foundation Child Health Research Awards Program (H.M.). Wolfgang Von Liedkle (Duke University) provided TRPV4-floxed mice. Hartmut Weiler (Blood Center of Wisconsin and Medical College of Wisconsin provided PAR1^-/-^ mice.

## AUTHOR CONTRIBUTIONS

D.L.W. conceived, designed and performed experiments and drafted the manuscript. S.E., N.N.J. prepared bronchial biopsies. R.B.S. and J.E. G. provided ciliated cells grown in air-liquid interface. M.B.F. and J.M.V. prepared and supplied club cells. D.J., M.E., and C.O. collected data from COAST patients and analyzed the data and assisted in writing the manuscript. T.W. analyzed histology and bronchial tissue biopsies. R.M. and G.K. assisted with analysis of *TRPV4* genotype and SNP analysis. B.S.K assisted with the study concept, design, and execution and helped write the manuscript.

## DELCARATION OF INTERESTS

The authors declare no competing interests.

## STAR METHODS

### LEAD CONTACT AND MATERIALS AVAILABILITY

Further information and requests for resources and reagents should be directed to and fulfilled by lead contact, Bruce S. Klein.

### EXPERIMENTAL MODEL AND SUBJECT DETAILS

#### Mice

All mice were C56BL/6 genetic background and were used for experimentation between the ages of 8 and 16 weeks. Males and females were used throughout the study. Sex and age had no obvious confounding effect in this model, but animals were age and sex matched for scientific rigor, nonetheless. Mice were bred in house and maintained in specific pathogen free conditions. All procedures were approved the Institutional Care and Use Committee. See **Key Resource Table** for an inventory of the transgenic and knockout animals used in this study.

### Human

Paraffin-embedded bronchial biopsies were obtained by bronchoscopy from subjects with asthma and from normal subjects as previously described (Nakamura et al., 2004). The study was approved by the University of Wisconsin-Madison Human Subjects Committee. Human lung epithelial cells were obtained from the tracheas of donor lungs and were differentiated at an air liquid interface (ALI) as described (Ashraf et al., 2015). Human adult airway basal stem cells were also recovered from discarded lung tissue obtained from the New England Organ Bank from donors under IRB approved protocols. These cells were isolated and differentiated into club and goblet cells as described (Feldman et al., 2019; Mou et al., 2016)

Participants of the COAST birth cohort study (initial *N*=289) were recruited in Madison, Wisconsin and surrounding areas from November 1998 to May 2000 (Lemanske, 2002). The study was approved by the University of Wisconsin-Madison Human Subjects Committee, and all families provided informed consent before enrollment. When the children reached 7 years of age, they themselves provided assent. Asthma was defined by parental report of physician diagnosis and/or use of asthma medications as previously described (Jackson et al., 2008). All COAST children were genotyped with the Illumina Infinium Exome Core array, and untyped SNPs (including rs6606743) were imputed using 1000Genome reference panel (CEU) as the reference.

### METHOD DETAILS

#### Allergic Inflammation Model

Mice were sedated with isoflurane. 25μg of *Aspergillus* protease or heat-inactivated (95°C, 10 minutes) control protease in 25μL of PBS was placed into the mouse pharynx. The tongue of the mouse was retracted with a forceps until the solution was completely aspirated into the lungs. This procedure was repeated on days 1, 2, 7, and 14 post-initiation. Lungs were harvested 24 hours after the last dose and processed for histology or flow cytometry. The protocol was also modified to track antigen-specific T cells. 10^6^ splenocytes from CD90.1+ OTII mice were injected into the tail vein 24 hours prior to the initial protease treatment. 25μg of ovalbumin was added to the protease inoculum, and all other aspects of the treatment remained unchanged.

#### Histology

Animals were euthanized by CO_2_ asphyxiation. The lungs were inflated with 10% paraformaldehyde via the trachea, the trachea was sutured shut, and the lungs were removed and placed in 10% paraformaldehyde for 1 hour. For histological dyes, the fixed lungs were rinsed with PBS, embedded in paraffin, cut into 10μm thick sections, and stained with periodic acid-Schiff or hematoxylin and eosin dyes. For immunofluorescence, the lungs were transferred to cryoprotectant (30% sucrose) and kept at 4°C for ∼4 days or until the lungs fell to the bottom of tube (i.e. fully saturated). The preserved lungs were washed with PBS, embedded in optimal cutting temperature (OCT) compound, cut into 8μm sections, and stored at −80°C. The slides were retrieved from −80°C storage, warmed to room temperature, soaked in PBS to remove OCT, and air dried for 15 minutes. Slides were incubated with Animal-free Block and Diluent (AFBD) with anti-mouse Fab fragment antibodies for 1 hour at room temperature in a humidified chamber. Slides were washed three times with PBS before adding primary antibodies at the indicated concentrations (see **Key Resource Table**) in AFBD. The slides were stored overnight at 4°C. The next day, the slides were washed three times with PBS, and the respective fluorophore-conjugated secondary antibody was added to the slides for 1 hour at room temperature in a humidified chamber. The slides were washed once with PBS, and 1μg/mL of DAPI was added to the slide for 10 minutes at room temperature. The slides were washed three times and allowed to air dry before adding hardset antifade reagent and coverslips. ∼24 hours later, a Nikon A1R confocal microscope was used to capture images of the slides. Images were processed with Fiji (Schindelin et al., 2012). For immunocytochemistry, we followed the manufacturer’s instructions without modification. TRPV4 staining intensity in the bronchial biopsies was scored by a blinded, surgical pathologist on a scale from 0 (weak) to 3 (strong).

#### Cellular Analysis

3μg of anti-CD45 APC-e780 antibodies were injected into the tail veins of mice to mark cells circulating in the blood vasculature. After 3 minutes, mice were euthanized by cervical dislocation. The lungs were harvested and minced with a gentleMACS (Miltenyi, San Diego, CA, USA) in digest solution, containing: RPMI-1640, 5% fetal bovine serum, 1500 U/mL type-1 collagenase, 1 mM CaCl_2_, 1 mM MgCl_2_. The samples were agitated at 37°C for 1 hour and further disrupted by gentleMACS. The cell solution was passed through a 70 μm filter, pelleted, and resuspended in 44% Percoll-RPMI medium. A Percoll density gradient was created (40% top, 67% bottom), and the samples were centrifuged for 20 min at 650 x g. The leukocytes at the interface were removed, washed twice with PBS + 0.1% bovine serum albumin (i.e. FACS buffer). ∼1/4 of each sample was placed in a 96 well plate. 1:500 LIVE/DEAD Fixable Dye and Fc block (1:100) was added to the samples for 10 minutes. The sample was washed and incubated for 1 hour at 4°C with fluorophore-conjugated primary antibodies (see **Key Resource Table**). The samples were washed, resuspended in FACS buffer, and processed by flow cytometry (LSRFortessa, BD Bioscience, San Diego, CA, USA). 50,000 count beads were added to the sample immediately prior to data collection. The data were analyzed with Flowjo X. See **Supplementary Figure 2** for gating strategy.

#### Mouse Treatments

For IL-5 blockade and CD4+ T cell depletions, mice were injected into the peritoneal (IP) cavity with 1 mg of antibody on days −1, 6, and 13 post-inoculation with protease. For phagocyte depletion, mice were given 3 intranasal doses of clodronate liposomes, followed by a 24-hour rest period before protease inoculation. To induce cre activity in Scb1a1-creERT2 mice, mice were injected IP with 2mg of tamoxifen on days −7 to −4, −1, 6, and 13 days post protease inoculation. For AAV6 *in vivo* gene expression experiments, mice were infected with 10^11^ AAV6 weekly for 3 consecutive weeks before commencing protease treatment. For junction targeting experiments, 25μg of poly L-arginine or poly L-glutamic acid were aspirated by mice on the same schedule as protease treatments. For thrombin inhibition experiments, mice were given 5U hirudin in the normal protease solution.

#### *In vivo* Antigen Capture/Presentation Assay

100μg of Eα-RFP was instilled into the lungs of sedated mice. 6 hours later, the mice were euthanized. The lungs were processed for flow cytometry, as explained above.

#### Calcium Flux Flow Cytometry

After euthanasia, 1ml of dispase was injected into the lungs of tamoxifen-treated Scgb1a1-creERT2 x LSL-Cas9 GFP mice. The lungs were removed, placed into collagenase solution, minced with a gentleMACS, and incubated with agitation for 30 minutes at 37°C. The lung cell suspensions were loaded with calcium dye (1μM Indo-1, 1mM MgCl_2_, 0.1% pluronic acid, 4mM probenecid) for 30 minutes at 37C. The samples were washed, stained with fluorescent antibodies, and finally suspended in calcium (1mM CaCl_2_) containing running buffer. 15 seconds of events were collected on the cytometer before retrieving the sample, adding 100μM final concentration of TRPV4 agonist, and replacing the sample in the cytometry for additional data collection.

#### Cytokine

Lungs from mice 0, 3 or 15-days post-inoculation were excised, snap frozen in liquid nitrogen, and homogenized in 2 mL of Tissue Protein Extraction Reagent with Complete Protease Inhibitor Cocktail. The lung homogenate was pelleted, and the supernatant was collected and stored at 80°C until analysis. CCL2 and IL-5 were measured in lung lysates by ELISA.

#### Co-immunoprecipitation

Club and goblet cells were generated as previously described (Feldman et al., 2019). ∼2×10^7^ cells collected from 2 T75 flasks flash frozen in liquid nitrogen. Membrane fractions were obtained (Sokabe et al., 2010), and the immunopreciptation was performed according to the manufacturer’s instructions.

#### *Ex Vivo* Restimulation

Lung leukocyte suspensions were incubated for 2 hours with 100μg of heat-inactivated protease in restimulation buffer (RPMI, 10% fetal bovine serum, 1X pen/strep, 1μM CaCl2, Beta-mercaptoethanol). 1X Brefeldin A was added to the well for the remaining 4 hours of incubation. At the conclusion, the cells were washed, stained with live/dead dye and surface antibodies, fixed and permeabilized with Foxp3 transcription buffer set, and incubated over night with antibodies to intracellular targets (i.e. Foxp3, IL-5, and GATA3).

#### Epithelium Permeability

125μg of Sulfo-NHS-Biotin in 25 uL of PBS + 1mM CaCl_2_ with protease or heat-inactive protease was aspirated into the lungs of sedated mice. After 1 hour, the lungs were processed for histology, as described above. Barrier integrity of primary human epithelial cells grown *ex vivo* was also evaluated with two methods. 2 mg/mL of 4kDa FITC-dextran with serial dilution of protease were added to the insert of the ALI device and left overnight in a tissue culture incubator. 100 μL of solution from the lower chamber of the ALI device was collected and 485/530 nm fluorescence was measured with a fluorimeter. Lastly, transepithelial electrical resistance was measured with a voltohmeter after 1 hour of incubation with varying concentrations of Alp1.

#### Enzyme Assays

All enzymatic reactions were performed at room temperature in 25mM Tris-base + 15mM NaCl + 10mM CaCl_2_, pH7.2. To assess cleavage activity, 2-fold serial dilutions of Alp1 were incubated with 2.5mg/mL FITC-in an opaque 96-well plate. 485/530nm Fluorescence intensity was detected after 15 minutes. 100μg human E-cadherin or 5μg C3 was incubated with 500ng of Alp1 for 1 hour, and western blots were performed with the reactions. 10mM of protease activated receptor peptides or 20mg/mL of human fibrinogen was incubated with 1mg/mL Alp1, 500U thrombin, or 10mg/mL trypsin for one hour. The solutions were analyzed by MALDI-TOF.

#### Quantification and statistical analysis

In the COAST patient cohort, associations between genotype and rates of allergic sensitization and asthma were assessed with an additive genetic model using the chi-square test for trend in proportions. All other statistical tests were Mann-Whitney U test with Bonferoni adjustment for multiple comparisons when >2 groups were compared. P<0.05 was deemed statistically significant.

#### Data and code availability

This study did not generate/analyze any large-scale datasets or computer code.

#### Abbreviations

Animal-free Block and Diluent (AFBD), Transepithelial Electrical Resistance (TEER), Air-liquid interface (ALI), Human Bronchiolar Epithelial Cells (HBEC), Transient receptor potential cation channel subfamily V member 4 (TRPV4), days post inoculation (dpi), Helper T (Th) cell, Innate lymphoid cell (ILC), Interleukin (IL), Monocyte-dendritic cell (Mo-DC), Eosinophil (Eos), C-C Chemokine Receptor (CCR), C-C Chemokine Ligand (CCL), Protease Activated Receptor (PAR), Toll-like Receptor (TLR), Calcineurin (CNB1), Scgb1a1 (Secretoglobin), Inhibitor of Nuclear Factor kappa-B Kinase Subunit beta (IKK), Nuclear Factor kappa-light-chain-enhancer of Activated B cells (NF-κB).

## SUPPLEMENTAL DATA

**Supplemental Figure 1. Recombinant *Aspergillus* alkaline protease 1. A)** Coomassie stained gel of culture supernate from *Pichia pastoris* transformed with Alp1 or untransformed. Alp1 is a ∼33kDa protein. **B)** Protease activity assay (FITC-Casein) of titrations of concentrate of culture supernate from Alp1 transformed (active/boiled) and untransformed *P. pastoris*. **Abbreviations**: Alp1 = alkaline protease 1. These are representative data from two or more independent experiments.

**Supplemental Figure 2.** Multiplexing antibodies to identify 17 pulmonary leukocyte subsets with 12 color flow cytometry. **A)** Leukocytes harvested from lungs of mice 15 dpi with Alp1 and stained with the following antibodies: FITC - Siglec F/TCRLδ; PercpCy5.5 - B220; PE - CD64/IL-33Rα; PECy7 - CD11c/TCRβ; BUV395 - Ly6G/CD4; BV421 - CD103/NK1.1; BV510 - Ly6C; BV650 - CD11b/CD8; BV785 - CD90.2; APC - CCR6; AF700 – MHCII; APCe780 - Dead/IV. **B)** Comparison of lymphocytes and myeloid cells analyzed by two methods. **C)** Cellular response in control and protease treated mice 15 dpi. **Abbreviations**: Alv Mac = Alveolar macrophages, CCR = C-C chemokine receptor, cDC = conventional dendritic cell, Eos = Eosinophils, IL = Interleukin, ILC = Innate Lymphoid Cell, Neu = MAIT = Mucosal-associated invariant T cells, Mo-DC = monocyte-dendritic cells, NK = Natural killer, TCR = T cell receptor, Th = Helper T cell. These are representative data from two or more independent experiments.

**Supplemental Figure 3. CCR2, but not CCL2, is required for allergic inflammation.** Mo-DC, IL-33R+ Th cell, and eosinophil response to Alp1 by **A)** CCR2^-/-^ or **B)** CCL2^-/-^ mice 15 dpi. **C)** Alveolar macrophages from the lungs of clodrosome- and control-treated mice. **D)** Confocal images of lung sections from mice 6 hours post-protease treatment. Green dye indicates CCL2 expression and red dye indicates epithelial cell subset (acetylated tubulin = ciliated cell; surfactant protein C = type-2 alveolar cell; CGRP = pulmonary neuroendocrine cell). 100X objective. **Statistics**: unpaired, non-parametric T test; * *P* < 0.05. **Abbreviations**: Alv. Mac. = Alveolar macrophage, CCL = C-C chemokine ligand, CCR = C-C chemokine receptor, CGRP = Calcitonin gene-related peptide, dpi = days post-inoculation, Eos = Eosinophils, IL = Interleukin, Mo-DC = monocyte-dendritic cells, Th = T Helper cell. These are representative data from two or more independent experiments.

**Supplemental Figure 4. Protease activated receptors 1 and 2 do not detect Alp1. A)** A peptide surrogate for the cleavage domain of PAR1 was incubated with fungal protease or thrombin control, and the cleavage products were measured by mass spectrometry. 1=RTDATVNPR; 2=SFFLRNPSENTFELVPLGDEE; 3=LRNPSENTFELVP or RNPSENTFELVPL; 4=LRNPSENTFELVPL; 5=FLRNPSENTFELVPL. **B)** Same as A) with PAR2 peptide. Mo-DC, Th2 cells, and eosinophils from lung of naïve or protease-treated wild type and **C)** PAR1^-/-^ or **D)** PAR2^-/-^ mice. **Statistics**: unpaired, non-parametric T test; * *P* < 0.05. **Abbreviations**: dpi = days post-inoculation, Eos = Eosinophils, IL = Interleukin, Mo-DC = monocyte-dendritic cells, PAR = Protease activated receptor, Th = T Helper cell. These are representative data from two or more independent experiments.

**Supplemental Figure 5. Alkaline protease 1 does not cause inflammation via release of C3a from Complement C3. A)** C3aR expression by club cells (Scgb1a1+) in naive mice**. B)** Western blot of purified C3a, C3, and C3 incubated with Alp1. **C)** MALDI-TOF analysis of B). **D)** Cellular immune response in the lungs of wild type or C3^-/-^ mice 15 dpi. **Statistics**: unpaired, non-parametric T test; * *P* < 0.05. **Abbreviations**: C = complement, dpi = days post-inoculation, Eos = Eosinophils, IL = Interleukin, Mo-DC = monocyte-dendritic cells, Th = T Helper cell. These are representative data from two or more independent experiments.

**Supplemental Figure 6. Fungal protease does not interact with the coagulation pathway. A)** Cellular response 3 dpi in mice treated with PBS, thrombin, protease, or thrombin + protease. **B)** Mo-DC and eosinophils from lungs of mice 3 days post treatment with PBS or protease in the presence and absence of a thrombin inhibitor (hirudin). **C)** Human fibrinogen incubated for 15 minutes with thrombin or Alp1. Samples were tilted to assess clotting of the solution**. D)** Mass spectrometry analysis of cleavage products (fibrinopeptide A or B) from clotting assay in panel C. **E)** Mo-DC and Eosinophils from the lungs of mice treated with fibrinogen, fibrinogen + heat-inactivated protease, or fibrinogen + functional protease**. F)** Cellular response to Alp1 15 dpi in the indicated knockout mice. **Statistics**: unpaired, Mann-Whitney U test with Bonferroni adjustment for multiple comparisons; * *P* < 0.05. **Abbreviations**: dpi = days post-inoculation, Eos = Eosinophils, IL = Interleukin, ILC = Innate lymphoid cells, F+P+H = Fibrinogen + Protease + Heat-inactivation, Mo-DC = monocyte-dendritic cells, Th = T Helper cell, TLR = toll-like receptor. These are representative data from two or more independent experiments.

**Supplemental Figure 7. Calcineurin and IKK2 are dispensable in type-2 alveolar cells, pulmonary neuroendocrine cells, and ciliated cells for the allergic response to protease.** Mice were challenged with Alp1 and lungs harvested 15 dpi. The cellular immune respone was quantified in mice with epithelial subset-specific deletion of **A)** calcineurin or **B)** IKK2. **Statistics**: One-way ANOVA. **Abbreviations**: Ascl1 = Achaete-scute homolog 1, dpi = days post-inoculation, Foxj1 = forkhead box J1, Mo-DC = monocyte-dendritic cells, Scgb1a1 = Secretoglobin, Th = T Helper. Compiled data from at least two independent experiments.

