## Supplementary figures and images for "Club cell TRPV4 as a damage sensor driving lung allergic inflammation"

### Supplemental Figures

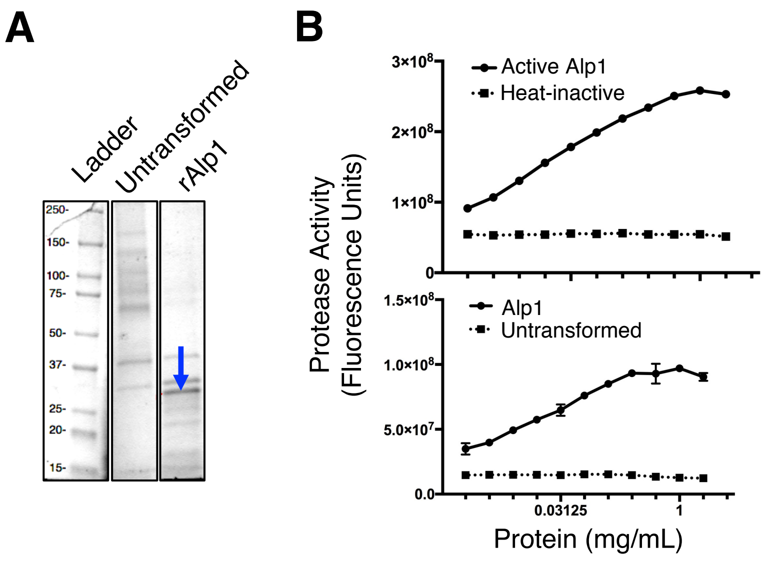

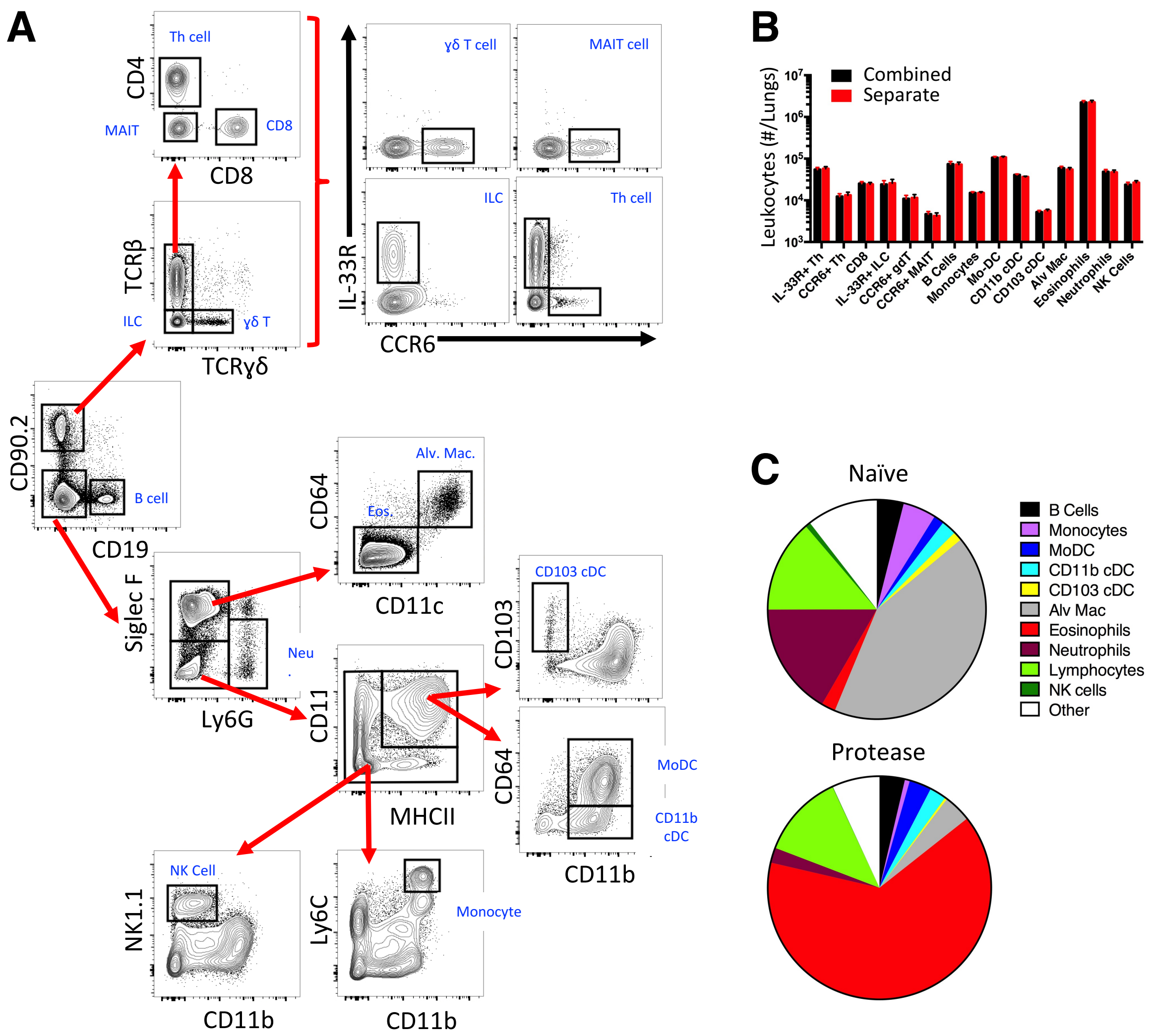

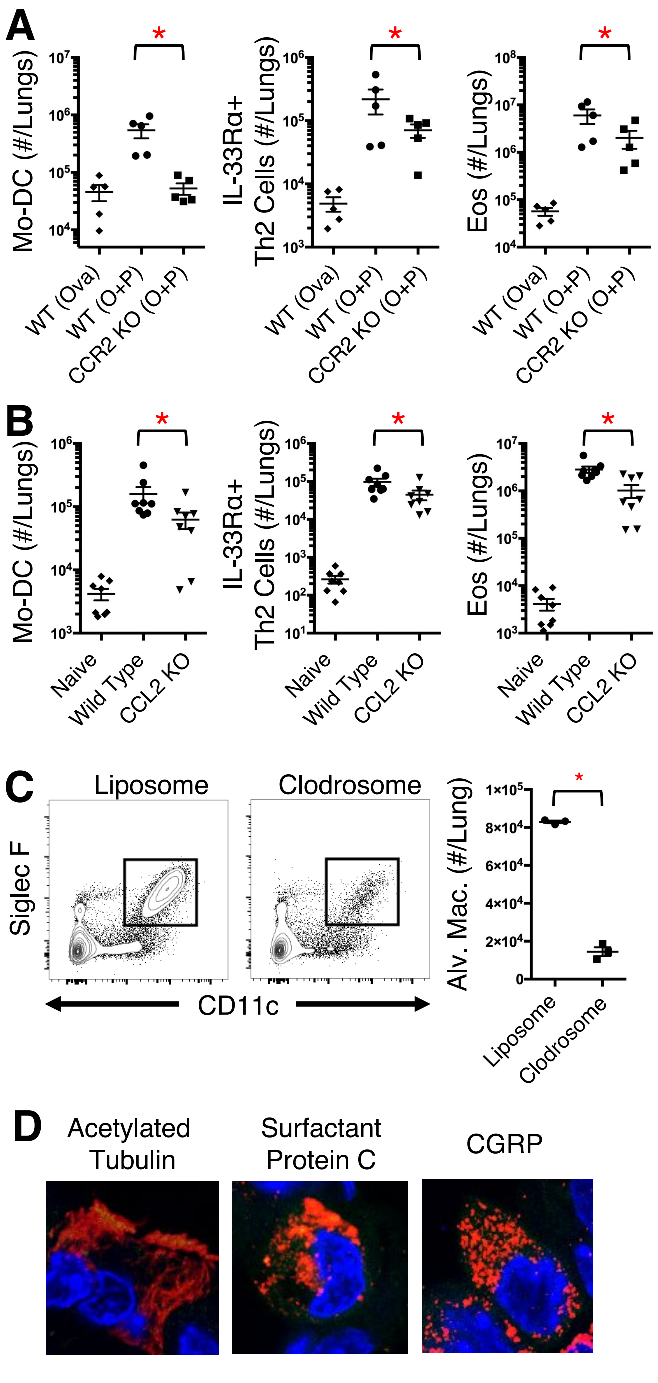

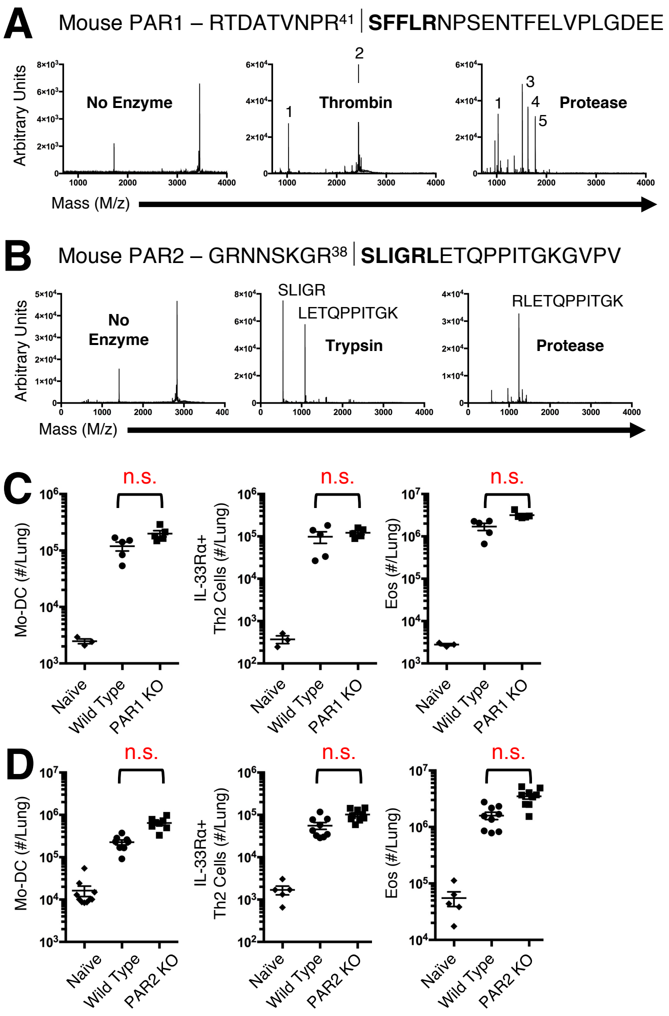

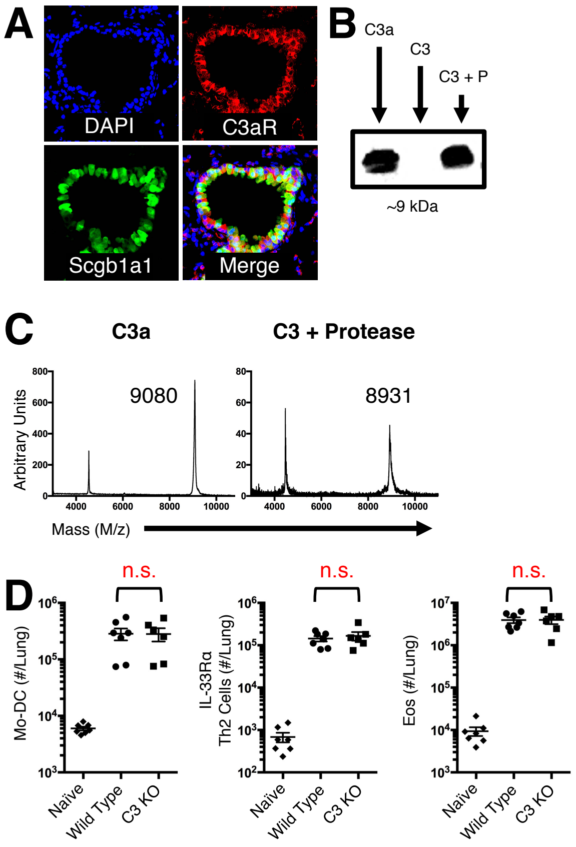

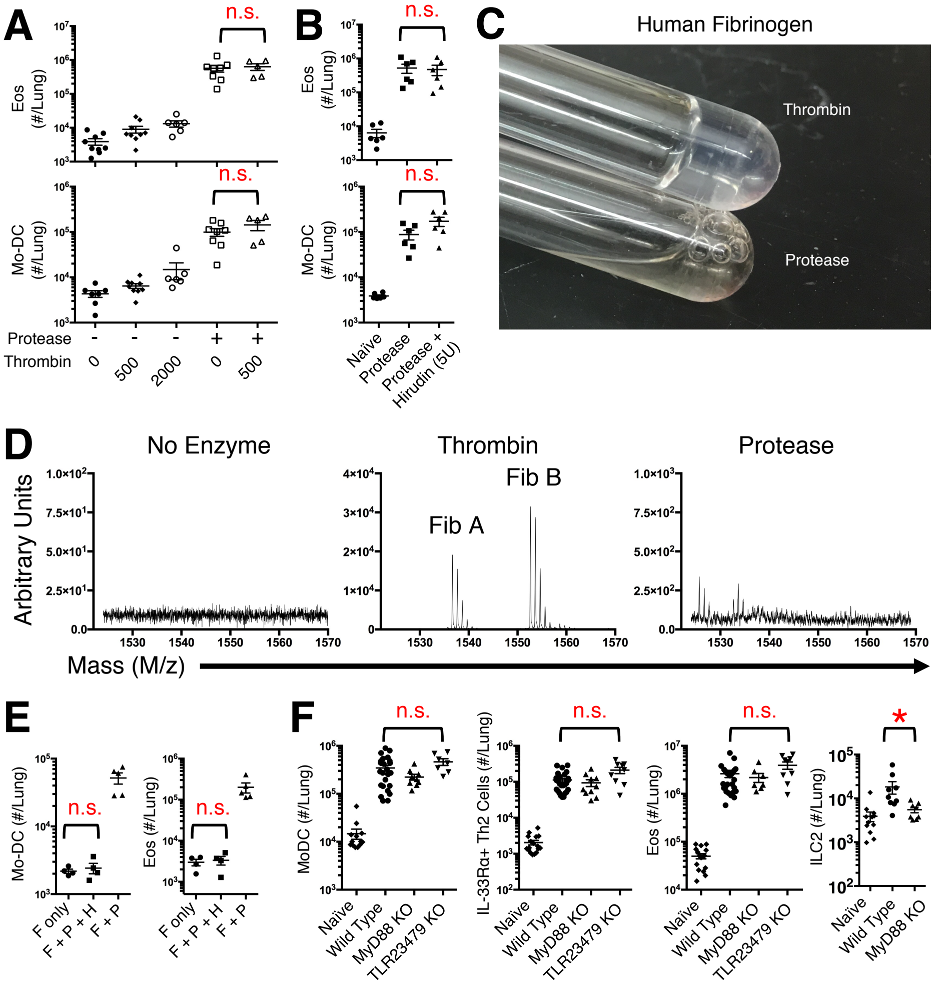

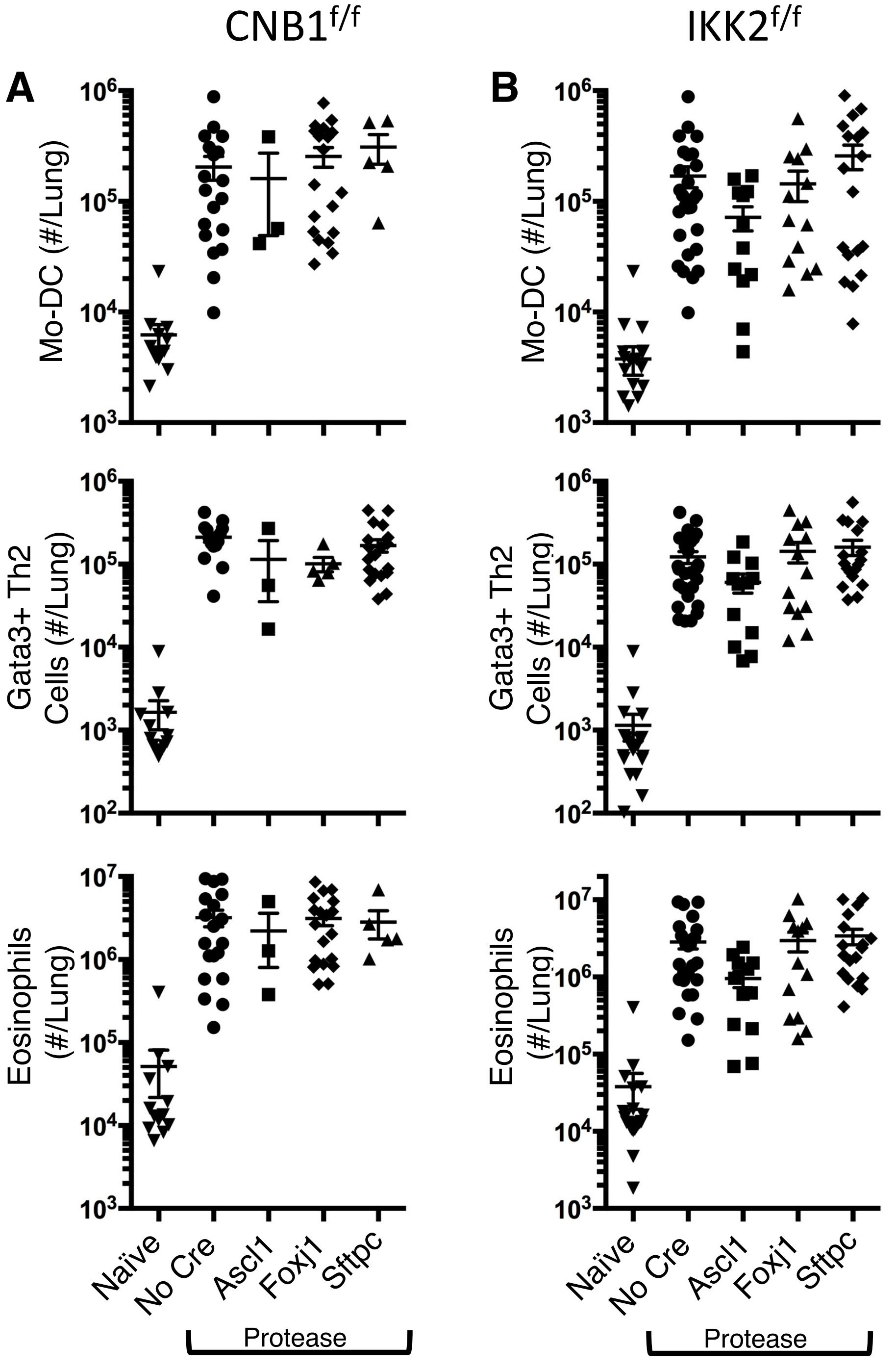
